# A Single Trophoblast Layer Acts as the Gatekeeper at the Endothelial-Hematopoietic Crossroad in the Placenta

**DOI:** 10.1101/2024.07.12.603303

**Authors:** Pratik Home, Ananya Ghosh, Ram Parikshan Kumar, Soma Ray, Sumedha Gunewardena, Rajnish Kumar, Purbasa Dasgupta, Namrata Roy, Abhik Saha, Madhu M. Ouseph, Gustavo W. Leone, Soumen Paul

**Author notes:** Corresponding authors. Correspondence: Soumen Paul, Pratik Home, Ram Kumar. Equal contribution (Pratik Home, Ananya Ghosh, Ram Parikshan Kumar).

## Abstract

During embryonic development the placental vasculature acts as a major hematopoietic niche, where endothelial to hematopoietic transition ensures emergence of hematopoietic stem cells (HSCs). However, the molecular mechanisms that regulate the placental hematoendothelial niche are poorly understood. Using a parietal trophoblast giant cell (TGC)-specific knockout mouse model and single-cell RNA-sequencing, we show that the paracrine factors secreted by the TGCs are critical in the development of this niche. Disruptions in the TGC-specific paracrine signaling leads to the loss of HSC population and the concomitant expansion of a KDR+/DLL4+/PROM1+ hematoendothelial cell-population in the placenta. Combining single-cell transcriptomics and receptor-ligand pair analyses, we also define the parietal TGC- dependent paracrine signaling network and identify Integrin signaling as a fundamental regulator of this process. Our study elucidates novel mechanisms by which non-autonomous signaling from the primary parietal TGCs maintain the delicate placental hematopoietic- angiogenic balance and ensures embryonic and extraembryonic development.

## Introduction

Trophoblast cells of the placenta establish a vascular connection between the mother and the fetus and express hormones that are essential for the successful progression of pregnancy. Placenta also acts as one of the major organs for hematopoietic stem cell generation, and the mid-gestation mouse placenta plays a significant role in the HSC development where it provides a temporary niche for definitive HSC pool ^1–3^. Defective development of placental hematopoiesis and vasculogenesis leads to serious pathological conditions such as preeclampsia and intra uterine growth restriction (IUGR)/ fetal growth restriction (FGR). These disorders result in pregnancy-related complications, maternal, prenatal and neonatal mortality, and affect ∼2–8% of pregnant women worldwide ^4^. The pathogenesis in preeclamptic patients is believed to be a response of vasculature to abnormal placentation ^5^. Thus, to define therapeutic modalities against these pregnancy-associated disorders, it is crucial to understand the molecular mechanisms that are associated with the proper development of placental hematopoiesis and vasculogenesis.

Trophoblast cell differentiation in the placenta involves mechanisms by which secreted paracrine factors within the placenta, and from the fetus and the mother regulate embryonic and extraembryonic development ^6^. Embryonic hematopoietic sites are characterized by the interlinked developments of the vascular and hematopoietic systems. Several studies have shown that the hemangioblasts and hemogenic endothelium act as the presumptive precursors to emerging hematopoietic cells ^7^. It has been indicated that the definitive hematopoiesis is autonomously initiated in the placenta, which subsequently generates HSCs from hemogenic endothelium and provides a niche for expansion of aorta-derived HSCs ^3,8^. Moreover, secreted pro-angiogenic factor, like Placental Growth Factor (PLGF), a member of the Vascular Endothelial Growth Factor (VEGF) family, and anti-angiogenic factors, like soluble Vascular Endothelial Growth Factor receptor-1 (sFLT1) and Endoglin (ENG), are expressed in the mouse placenta, implying autocrine or paracrine actions ^9,10^. How these signaling mechanisms influence the development of the maternal-fetal interface are poorly understood.

Several studies have implicated GATA family of transcription factors in the development of HSCs in other organs, and previously we have shown that GATA2 and GATA3 are involved in the trophoblast development and differentiation ^11–15^. They are implicated in the regulation of the expression of several trophoblast-specific genes, including prolactin hormone Placental lactogen I (*Prl3d1*, also known as *Pl1*) and the pro-angiogenic factor Proliferin (*Prl2c2*, also known as *Plf*) ^16–18^. Recently we demonstrated that placenta-specific redundant functions of *Gata2* and *Gata3* are important in trophoblast lineage development ^19^. The simultaneous knockout of *Gata2* and *Gata3* resulted in significant developmental defects in the placenta as well as the embryo proper, leading to very early embryonic lethality ^19^. These developmental defects were accompanied by severe blood loss in the placenta, yolk sac, and the embryo proper. Moreover, we established dual conditional *Gata2* and *Gata3* knockout trophoblast stem cells and used ChIP-Seq and RNA-Seq analyses to define independent and shared global targets of GATA2 and GATA3. We found that several pathways associated with the embryonic hematopoiesis and angiogenesis are targets for both transcription factors^19^. However, very little is known about how these two master regulators operate in a spatiotemporal manner within the mature trophoblast during placental development and dictate placental hematopoiesis and angiogenesis.

In a mouse conceptus, parietal trophoblast giant cells (parietal TGCs), which line the border of the growing placenta and the maternal decidua, is thought to be a major source of endocrine and paracrine signals during mouse placentation ^20,21^. Parietal TGCs are initially developed from the trophoectoderm (primary parietal TGCs) and subsequently from the secondary differentiation of precursors in the ectoplacental cone (secondary parietal TGCs). Gene expression analyses reveled that both the primary and secondary parietal-TGCs specifically express prolactin 3d1 gene (*Prl3d1*), also known as placental lactogen 1 (PL-1). Therefore, to understand how GATA2/GATA3 functions could dictate endocrine and paracrine functions in a developing placenta, we specifically deleted both *Gata2* and *Gata3* in the parietal TGC using *Prl3d1tm1(cre)Gle* (*Pl1-Cre*) mouse line ^20,22,23^ that drive Cre expression specifically within the parietal TGCs. We noticed in-utero death of *Gata2/3* conditional double knockout *Gata2^f/f^*;*Gata3^f/f^;Pl1^Cre/wt^* (GATA-Pl1 KO) embryos starting from embryonic day 12.5 (e12.5). A fraction of the GATA-Pl1 KO embryos survived to birth. However, majority of the surviving GATA-Pl1 KO pups showed severe growth restriction. Remarkably, we noticed that along with defect in trophoblast development, the delicate balance between hematopoietic vs angiogenic differentiation is lost in the developing GATA-Pl1 KO placentas. The GATA-Pl1 KO placentas showed severe defect in hematopoiesis due to reduced hematopoitic stem and progenitor cells and disorganized placental vasculature due to defective differentiation of endothelial progenitors. Thus, our study highlight the importance of a GATA-mediated transcriptional program within a single trophoblast subtypes that fine-tunes the hematopoietic and endothelial development during placentation.

## Results

### GATA deletion in the trophoblast giant cell layer in mouse embryos display embryonic and extraembryonic hematopoietic defects

Our previous studies have shown that the loss of GATA factors in the trophoblast lineage cells results in the gross phenotypic abnormality in the placenta and the embryo proper in a mouse model ^19^. The placenta contains distinct layers of differentiated trophoblast cells, each having specialized functions to support a pregnancy. These include TGCs, spongiotrophoblasts (SpT), glycogen trophoblasts, and labyrinthine trophoblasts. Each subclass is characterized by their unique gene expression signatures ^24^. Thus, it is imperative to analyze how GATA factors act in different subclasses of trophoblast cells in the context of placental development and function. As TGCs have been reported to express both GATA2 and GATA3, we chose to delete *Gata2* and *Gata3* together in the TGCs ^16^. Placental TGCs are marked by the expression of three prolactin family of proteins Prl2c2 (PLF), Prl3b1 (PL2), Prl3d1 (PL1) and Cathepsin Q (CTSQ)^20^. Out of these four genes, PL1 expression is restricted to the parietal TGCs only . Thus, to restrict GATA deletion exclusively to the parietal TGCs, we used a previously established *Prl3d1^tm1(cre)Gle^* (*Pl1^Cre^* ) mouse model ^22^ , where Cre recombinase is selectively expressed in the *Pl1* expressing cells (**Fig. S1A**). We validated the *Pl1*-specific Cre expression in this mouse model by LacZ staining (**Fig. S1B).** We also developed a murine model in which the parietal trophoblast giant cell layers were fluorescently labeled. We used *Gt(ROSA)26Sor^tm4(ACTB-tdTomato,-EGFP)Luo^/J*, (also known as *mT/mG*) mice, which possess loxP flanked membrane-targeted tdTomato (mT) cassette and express strong red fluorescence in all tissues ^25^. Upon breeding with the *Pl1*-Cre recombinase expressing mice, the resulting offsprings have the mT cassette deleted in the Cre expressing cells(s), allowing expression of the membrane-targeted EGFP (mG) cassette located in-frame immediately downstream. Microscopic analyses of the conceptuses and their cryosections from a cross between *Pl1^Cre^* male and *mT/mG* female confirmed the selective EGFP fluorescence only in the parietal TGC layers (**Fig**. **S1C**, **Fig. 1A, B**). This expression of *Pl1-Cre* is consistant with the recent publication which clearly illuminates the PL1 protein expression exclusively in the P-TGCs ^23^.

**Fig. 1:**
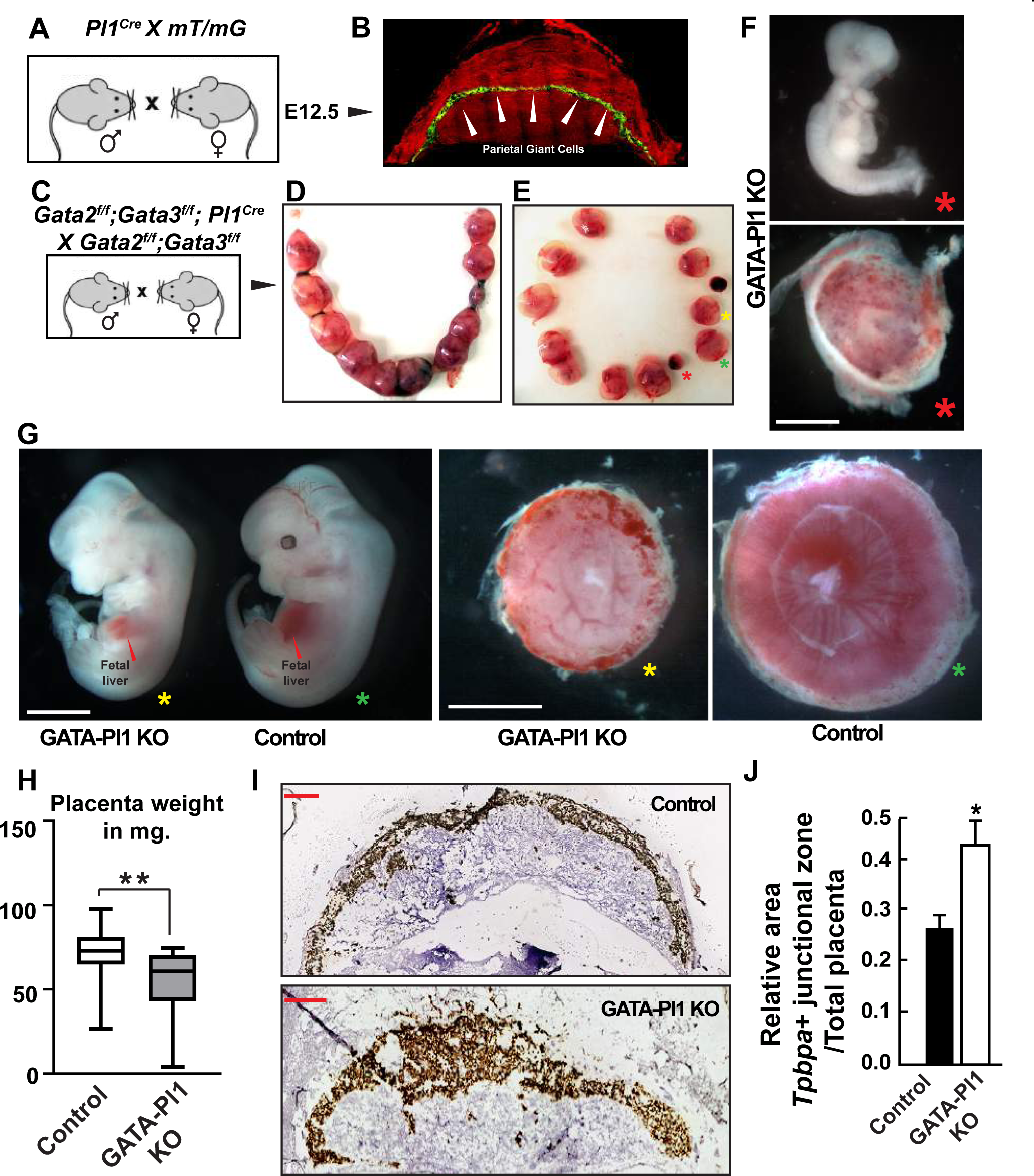
Trophoblast giant cell-specific GATA deletion leads to embryonic lethality and developmental defects. (A) Mating strategy to identify the Prl3d1 positive parietal giant cell layer. (B) Cryosections of the *mT/mG Pl1^Cre^* conceptus show exclusive EGFP expression in the parietal TGCs, indicating the specific nature of the *Pl1^Cre^* expression. (C) Mating strategy to define the importance of GATA factors in the trophoblast giant cells. (D) E12.5 uterine horn harvested from the above mating contains apparent embryonic resorption sites. (E) Isolated conceptuses associated with GATA Pl1-KO embryos show severe developmental defects (**\***) and size differences (**\*, \***) compared to the control (**\***). (F) Embryos isolated from one of the extremely small conceptuses (**\***) reveal gross developmental defects and embryonic death accompanied by small and thin placental tissue. Scale bar 2 mm. (G) Comparison between a GATA Pl1-KO embryo (**\***) and non-Pl1^Cre^ littermate (**\***) indicates fetal growth reduction and apparent blood loss , and defective vasculature in both the embryo proper and the placenta. Fetal liver (marked by red arrow) of the GATA Pl1-KO embryo also showed reduced hematopoiesis compared to the control. Scale bar 2 mm. (H) GATA Pl1-KO placentae shown in G (**\***) were of significantly smaller weight compared to the controls (**\***) (Mean±s.em., n=21 for the control, n=18 for the KO, **P≤0.01, analyzed by two-tailed Student’s t-test). (I) RNAScope labeling of implantation sites using the *Tpbpa* probe marks the spongiotrophoblast layers. Scale bar 1 mm (J) Quantitative comparison of the relative ratio of Tpbpa+ junctional zone area and labyrinth area between the control and GATA Pl1-KO implantation sites (Mean±s.em., n=5 for each samples, ***P≤0.001, analyzed by two-tailed Student’s t-test).

Using *Gata2^f/f^*;*Gata3^f/f^* mouse model reported in an earlier study from our laboratory, we established a conditional GATA knockout mouse model *Gata2^f/f^*;*Gata3^f/f^;Pl1^Cre/wt^* (GATA-Pl1 KO)^19^. Although a significant number of GATA-Pl1 KO embryos showed lethality and abnormal embryonic and extraembryonic phenotypes, a small percentage of them were viable. However, almost all of them showed visible growth restriction at birth (**Fig. S2**). These viable growth restricted pups were fertile and were chosen to function as breeders. For the embryo analyses, we used these males for mating with non-Cre females to restrict the effect of GATA deletion exclusively to the placental tissue. These mice were used to selectively knockout both *Gata2* and *Gata3* in the parietal trophoblast giant cells. PCR with *Gata2* and *Gata3* deletion specific primers confirmed the gene deletions specifically in the placenta in the GATA-Pl1 KO and not in the embryos (**Fig. S3A**). In addition, using laser captured microdissection (LCM) we excised GFP-positive pTGCs from *Gata2^f/f^*;*Gata3^f/f^;Pl1^Cre/wt^ mT/mG* placental sections. Again, PCR with *Gata2* and *Gata3* deletion specific primers confirmed efficient deletion of the *Gata2* and *Gata3* in these cells compared to the cells excised from adjacent labyrinth zone (**Fig. S3B**). Thus, we validated that *Gata2* and *Gata3* could be efficiently deleted in a highly pTGC-specific manner in the *Gata2^f/f^*;*Gata3^f/f^;Pl1^Cre/wt^* mouse model.

Previous studies have shown that the major expansion of the hematopoietic stem cell (HSC) population in the mouse placenta takes place between E11.5 and E13.5 ^1^. Thus *Gata2^f/f^*;*Gata3^f/f^; Pl1^Cre /wt^* males were crossed to *Gata2^f/f^*;*Gata3^f/f^* females and the embryos were analyzed between E10.5 and E13.5 (**Fig. 1C**). Resulting phenotypic abnormalities were observed mostly at E12.5 and E13.5. For all subsequent analyses, E12.5 and E13.5 conceptuses were chosen.

Two major groups of embryos with distinct phenotypic abnormalities were observed in the *Pl1^Cre^* positive embryos (**Fig. 1D, E**). One group showed very early embryonic death and was associated with extremely small fetal (necrotic) and placental tissues (**Fig. 1E, F**). The other group showed significant growth restriction, developmental defects, and blood loss in the embryo proper, while their placentae showed significant anomalies in size and weight and were associated with apparent blood loss (**Fig. 1E, G, H**). These placentae showed appernt loss of vasculature (**Fig. 1G**). These defects were most prominent at E12.5 and E13.5 (**Fig. S2**). Non-Cre embryos from the same littermates were treated as controls (**Fig. 1E, G**). Along with these gross abnormalities, significant alterations in the placental architecture were observed in the GATA-Pl1 KO samples. A marked increase in the junctional zone (marked by the spongiotrophoblast/ glycogen trophoblast marker *Tpbpa*) was observed compared to the control (**Fig. 1I-J)**.

Interestingly, the fetal liver in the GATA-Pl1 KO embryos showed blood loss (**Fig. 1G**), which is consistent with the hypotheses that along with the hematopoietic stem and progenitor cells (HSPCs) from the yolk sac, the placental HSCs migrate to and seeds fetal liver for hematopoiesis ^26^.

Thus, our study showed that the GATA factor-loss in the parietal TGCs of the placenta was sufficient to significantly impair embryonic and extraembryonic growth and also result in blood loss and vasculature defects.

### GATA factor functions in parietal trophoblast giant cells regulate trophoblast progenitor developments

Some of the critical tasks of TGCs are the secretion of autocrine and paracrine factors that are involved in the trophoblast outgrowth and placental development process ^27,28^. Genetic knockout models targeting TGC specific genes have been shown to affect lineage-specific trophoblast differentiation and thereby results in abnormal development of the placenta ^22,27,29,30^. As mid-gestation mouse placenta contains numerous cell types, including several different trophoblast subtypes, endothelial cells, hematopoietic cells (both HSCs, multi-lineage cells, and terminally differentiated cells), stromal cells, it is challenging to analyze the effect of a trophoblast-specific gene knockout on placental cell subpopulations. To analyze the effect of GATA deletion in the TGCs, we performed Single-cell RNA-Sequencing (scRNA-seq) analyses of E13.5 placenta from a pregnant *Gata2^f/f^*;*Gata3^f/f^* female crossed with *Gata2^f/f^*;*Gata3^f/f^; Pl1^Cre/wt^* male. Two individual GATA-Pl1 KO placentae and two individual control littermate placentae were used for the sequencing. The genotypes were confirmed by PCR using tissue from the embryo proper(genotype for GATA Pl1-KO sample 1 is shown in **Fig. S3A**). The knockout placentae where the apparent growth defects and blood loss phenotypes were most severe were chosen for the scRNA-seq analysis (**Fig. 1G**).

T-distributed Stochastic Neighbor Embedding (t-SNE) plot of the aggregated hierarchical clustering of the four samples revealed 33 distinct clusters (**Fig. 2A**). The clustering of both the control samples showed significant similarity to each other, while significant clustering similarities were observed in both the knockout samples, indicating a high degree of homology between the placentae of the same genotypes (**Fig. S4**). To categorize the clusters, we identified the top upregulated genes from each cluster (**Dataset 1**). Alongwith that, we used “single-cell Mouse Cell Atlas (scMCA) analysis” pipeline previously described in a single-cell RNAseq study on murine embryonic and extraembryonic tissues at E14.5 ^31^. We individually fed digital gene expression (DGE) matrix for each cluster to scMCA which in turn used a previously defined set of gene signatures ^31^ and predicted nature of that cluster with a probability score. Alongwith that, it also listed top markers associated with that cluster. Finally, we cross-matched the scMCA top markers from **Dataset 2** with the top markers we derived earlier (**Dataset 1**). In this way we broadly identified five different cluster types in our samples.

**Fig. 2:**
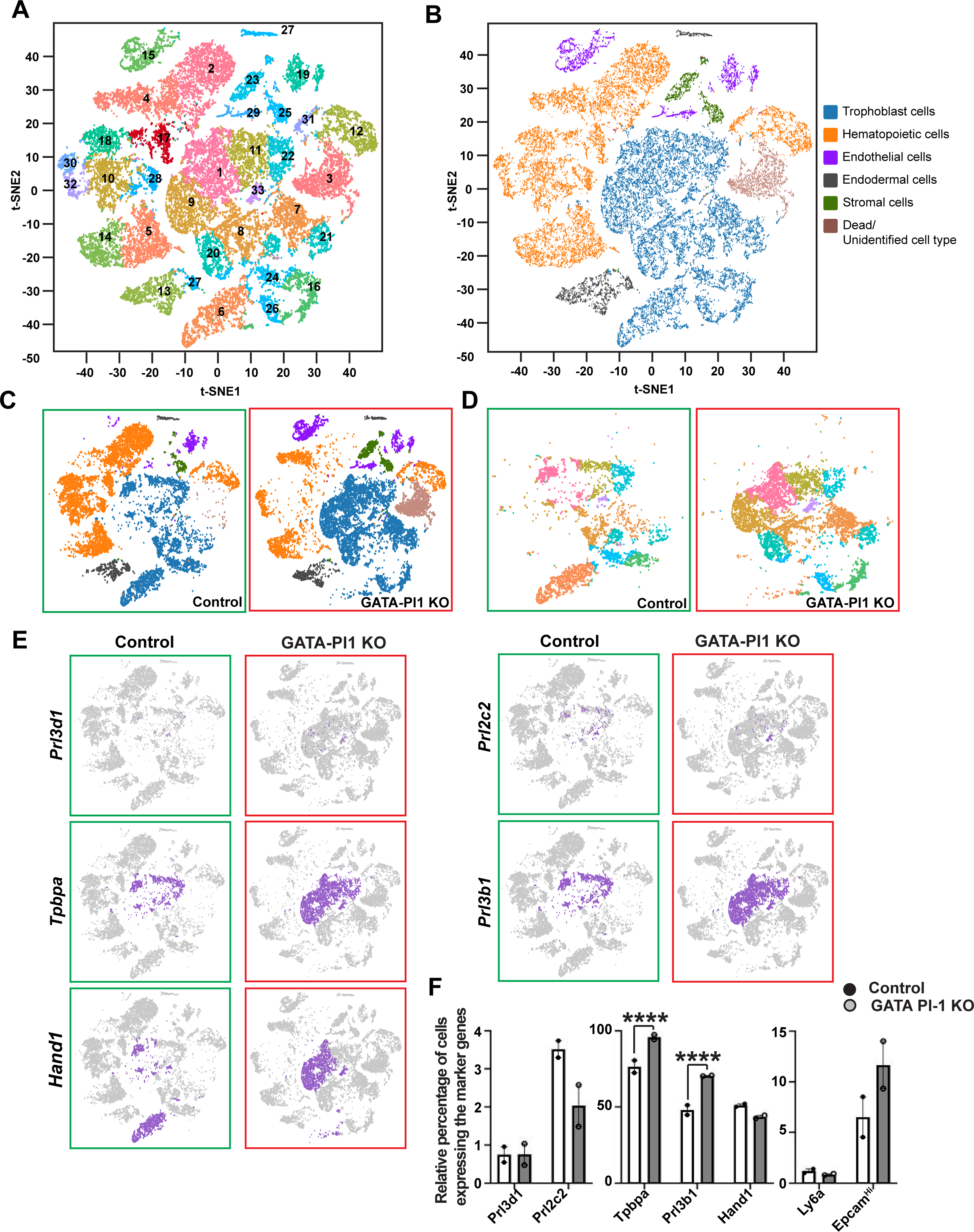
Single Cell RNA-Seq analyses of the TGC-specific GATA factor loss show altered placental trophoblast subpopulation. (A) A t-SNE plot of the aggregate of the hierarchical clustering of 2 control and 2 GATA Pl1-KO placental samples shows 33 distinct clusters. (B) t-SNE plots of the aggregate samples show major cell types. (C) Individual t-SNE plots of the control (aggregate of two control samples) and the KO (aggregate of two KO samples) samples show differences in the major cell subtypes. (D) Individual clusters representing trophoblast subtypes show major differences between the control and the KO placentae. Cluster 3, identified as dead cells/ unidentified cell type, were excluded. (E) Comparative t-SNE plot of cells marked by the expression (log 2-fold expression > 0, p-value <0.05) of trophoblast lineage markers *Prl3d1*, *Prl2c2*, *Tpbpa*, *Prl3b1*, *Hand1* in the control (green box) vs GATA Pl1-KO (red box) placentae. Expression in only the trophoblast cells are represented. (F) Quantitative analyses of the relative cell percentages for the *Prl3d1*, *Prl2c2*, *Tpbpa*, *Prl3b1, Hand1, Ly6a, and Epcam*^Hi^ positive cells. For all markers except *Epcam*^Hi^, log 2-fold expression > 0; for *Epcam*^Hi^ log 2-fold expression > 3. All cell numbers were normalized using total trophoblast cell numbers corresponding to the sample. (Mean±s.em., n=2 for each sample type, ****P≤0.0001, analyzed by two-tailed Student’s t-test).

They include trophoblast cells (clusters 1, 6, 7, 8, 9, 11, 16, 20, 21, 22, 24, 26, 33), hematopoietic cells (cluster 2, 4, 5, 10, 12, 14, 17, 18, 28, 30, 31, 32), endothelial cells (clusters 15, 19, 29), endodermal cells (clusters 13, 27), and stromal cells (clusters 23, 25) (**Fig. 2B, Dataset 2**). As scMCA could not predict the cell type for cluster 3, we looked at the top marker expression for this cluster. It showed significant enrichment of several mitochondrial genes (**Dataset 1**). Although contributions for the cluster 3 population came almost entirely from the GATA Pl1-KO samples (**Fig. 2C, S4**), the high level of mitochondrial gene enrichment in this cluster implied dead cells captured during the library preparation for scRNA-seq. Thus Cluster 3 was excluded for subsequent analyses. However, we can not exclude the possibility of cluster 3 being a yet unknown. Aggregates of two control samples and GATA-Pl1 KO placentae are used and represented as Control and GATA-Pl1 KO henceforth. Major alterations and rearrangement of the trophoblast cell populations, which was associated with significant loss of the hematopoietic cell populations, were evident in the GATA-Pl1 KO placentae compared to the control (**Fig. 2C**).

We also examined the enrichment of different trophoblast markers in individual trophoblast- clusters based on their log2-fold expression ratio of normalized mean gene UMI counts with a p-value < 0.05 ( **Dataset 2, 3**). Based on this analysis, we detected considerable gain in the spongiotrophoblast population, including invasive spongiotrophoblasts, clusters 1, 7, 9, and 11, which further validates our *in situ* hybridization observation above in **Fig. 1I-J** (**Fig. 2D**). We also observed a significant increase in the spiral artery-associated TGCs, clusters 8, 20, and 26 (**Fig. 2D**). This was accompanied by a significant loss in the high level of *Gjb3* expressing trophoblast progenitor population, clusters 6 and 24 (**Fig. 2D**). Cluster 16, which showed strong expression of syncytiotrophoblast markers *Dlx3*, *Tfeb*, *Hsd11b2*, gap junction protein *Gjb2*, PDGF receptor α (*Pdgfa*), *Pparg*, *Slc16a1*, did not show significant changes in the GATA-Pl1 KO placentae.

Although the scMCA analyses predicted cluster 33 to be of mixed nature including invasive spongiotrophoblasts, spiral artery trophoblast giant cells and progenitor trophoblasts (**Dataset 2**), we observed enrichment of several trophoblast giant cell associated genes in that cluster (**Dataset 1**). They include pregnancy specific glycoproteins (*Psg23*, *Psg18*, *Psg19*, ), several prolactin family members (*Prl8a9*, *Prl7d1*, *Prl3b1*), Trophoblast-specific protein alpha (*Tpbpa*), Endothelial protein C receptor (*Procr*), LIF receptor alpha (*Lifr*), Fibronectin 1 (*Fn1*). Moreover, along with strong *Tpbpa* expression, cluster 33 also contained cells positive for *Prl3d1*, *Prl2c2*, *Hand1*, *Prl3b1,* all markers for parietal trophoblast giant cells, spiral artery-associated trophoblast giant cells, canal trophoblast giant cells, and sinusoidal trophoblast giant cells ^32^ (**Fig. 2E**). Thus we identified cluster 33 as the TGC-cluster. Significantly, out of all the 33 clusters, only this cluster showed increased coexpression of *Prl3d1*, *Gata2* and *Gata3* (**Fig. S5)**. Interestingly, the loss of *Gata2* and *Gata3* in these cells did not seem to negatively affect the parietal TGC population (**Fig. 2E, F**).

A comparison between the control sample and the knockout sample populations showed a significant increase in the Trophoblast-specific protein alpha (*Tpbpa*) expressing cell population (**Fig. 2E, F**). These cells were also found to be positive for Prolactin family member *Prl3b1*. On the other hand labyrinth trophoblast progenitor cells marked by the expression of *Ly6a* (*Sca1*) and high levels of Epcam (*Epcam*^Hi^) ^33^ and *Hand1* expressing cell population did not show any significant alterations (**Fig. 2E, F**).

We also noticed defective syncytiotrophoblast (SynT) development in GATA-Pl1 KO placentae. The mouse labyrinth contains two layers of SynTs, namely SynT-I and SynT-II. which express two distinct moncarboxylate transporters (MCTs). The SynT-I expresses MCT1 [also known as Solute Carrier Family 16, Member 1 (SLC16A1)] and the SynT-II expresses MCT4 [also known as Solute Carrier Family 16, Member 3 (SLC16A3)]. Recent studies revealed that the labyrinth of a developing mouse placenta contains distinct progenitors for SynT-I and SynT-II layers^15,23^. The progenitors for SynT-I can be identified from the mRNA expression of *Glis1, Snap91, Stra6, Tfrc, Epha4* and *Slc16a1*, whereas the progenitors of SynT-II can be identified from the mRNA expression of *Igf1r, Egfr* and *Slc16a3*. From our scRNA-seq analyses, we noticed that the relative abundance of The SynT-I progenitors were not altered in E12.5 GATA-Pl1 KO placentae (**Fig. S6A**). However, a significant increase in SynT-II progeniors was noticed in GATA-Pl1 KO placentae. We also noticed increased number of *Gcm1* expression is gradually suppressed in SynTs after E9.5 and by E12.5 only a few SynT-II cells express *Gcm1* (**Fig. S6B**). Thus, increased abundance of SynT-II precursors and *Gcm1* expressing cells indicate an skewed SynT differentiation program in GATA-Pl1 KO placentae. Together, these findings revealed that the parietal TGC-specific loss of GATA factors skews the trophoblast differentiation process and thereby alters the distribution of trophoblast subpopulations in the developing placenta and affects gross placental architecture. Remarkably, these data also indicate a novel mechanism whereby the parietal TGC-specific paracrine signaling dictates the differentiation of the different trophoblast progenitors and regulates the development of distinct placental layers.

### Loss of GATA factors disrupts hematopoietic-endothelial cell lineage segregation and affects fetal hematopoiesis

The placenta is one of the major sites for *de novo* hematopoiesis in an embryo ^1^. It not only supports the expansion of the nascent HSC population but also protects the HSCs from premature differentiation cues. Several studies have implicated primary and secondary trophoblast giant cells in secreting paracrine and endocrine factors ^34^, which are thought to regulate placental hematopoiesis. These include prolactin/ placental lactogen class of hormones ^35^, interferon ^28^, vasodilators ^36^, anticoagulants ^37^ and several angiogenic factors ^29,38,39^. In order to analyze the effect of PL1-specific GATA factor loss on the placental hematopoietic cell development, we used our scRNA-seq data. Ingenuity Pathway Analysis (IPA) was performed to compare the physiological functions of the *Prl3d1+* cells between the control and the KO samples. Our results indicated that major physiological functions related to the hematopoiesis and angiogenesis were downregulated in the *Prl3d1+* cells in the GATA-Pl1 KO samples (**Fig. S7**).

We identified major hematopoietic subpopulations in our samples that include B-cells, T-cells, Granulocytes, Macrophages, Erythroblasts, Dendritic cells, Basophils, Megakaryocytes, Macrophages, and Natural Killer cells (**Fig. 3A, Dataset 2)**. Placental HSCs are localized in the labyrinth, and umbilical blood vessels ^40^ and have been shown to express surface markers KIT, CD34, and Ly6A ^2^. We found two major clusters which harbored *Kit*^+^ *Cd34*^+^ *Ly6a*^+^ cells, cluster 15 and cluster 17 (**Fig. 3A, B**). We calculated the relative percentages of cluster 15 and 17 cells with respect to the total cell numbers per sample. While cells in the cluster 17 show gross reduction in the GATA Pl1-KO, cluster 15 displayed opposite trend. Almost all cells in cluster 15 were the contribution from the GATA Pl1-KO placentae (**Fig. 3C)**. Moreover, when we calculated the relative percentages of the *Kit*^+^ *Cd34*^+^ *Ly6a*^+^ HSC population, we found marked reduction of these cells in the GATA Pl1-KO compared to the control (**Fig. 3D)**. Interestingly, unlike 17, cluster 15 is marked by high level of expression of vascular endothelial growth factor receptor- 2 (*Vgfr2/Kdr*) (**Fig. 3E**).

**Fig. 3.**
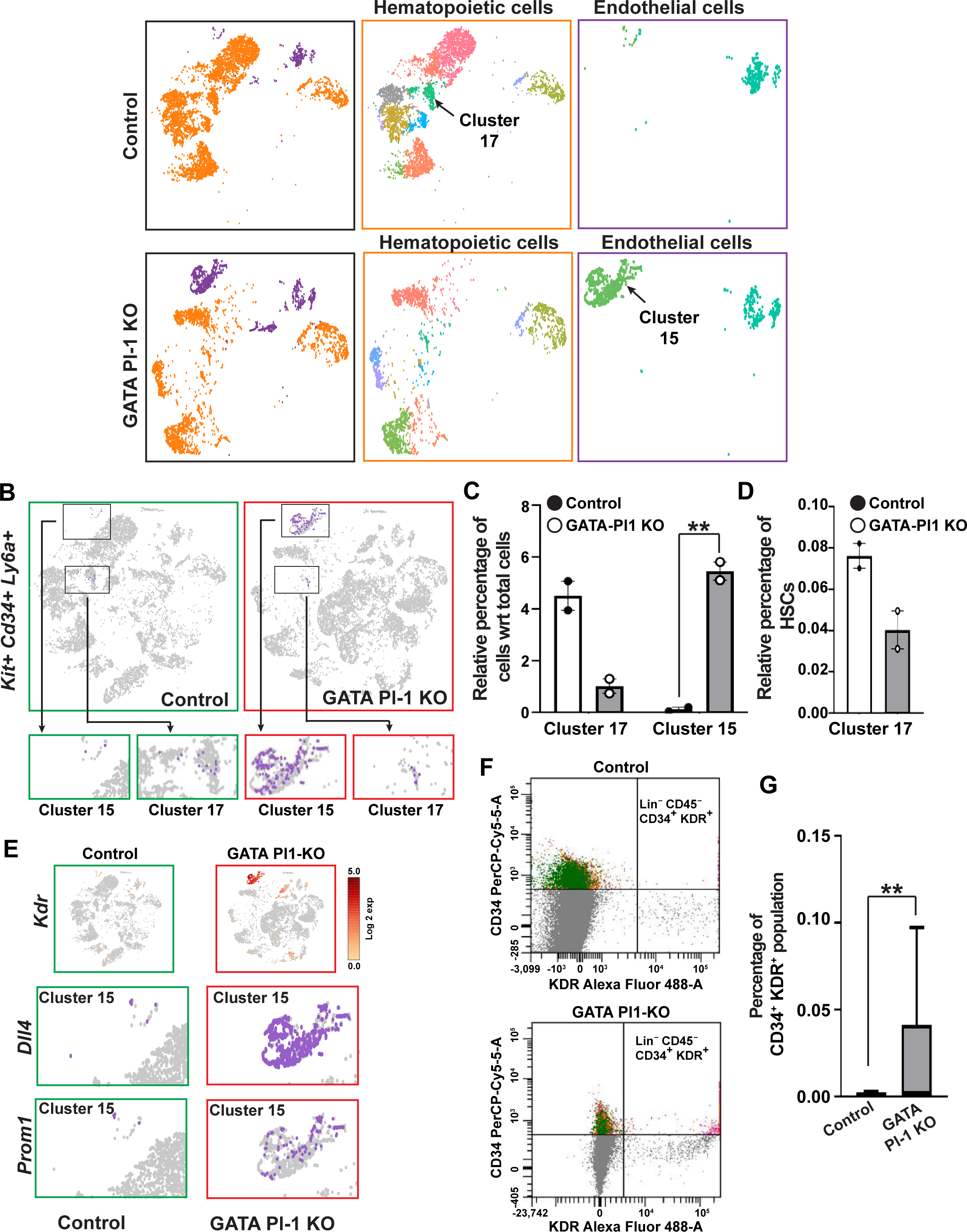
TGC-specific GATA factor function is essential for the differentiation of the hematoendothelial niche. (A) Comparative t-SNE plots representing the hematopoietic (orange) and endothelial (purple) cell populations in control vs. GATA Pl1-KO placentae are shown. Further cluster distribution of these two populations are shown with the orange box (hematopoietic cells) and the purple box (endothelial cells). These subclusters are described in Fig. 2. (B, C) scRNA-seq analyses show cluster 15 and 17 to harbor the *Kit*+ *Cd34*+ *Ly6A*+ population of HSCs. The t-SNE plots show a marked reduction in the overall cell numbers in cluster 17 and huge increase in the cluster 15 of the GATA Pl1-KO placentae compared to the control (Mean±s.e.m., n=2 for each sample type, **P≤0.01, analyzed by unpaired Student’s t-test, Two-stage step-up (Benjamini, Krieger, and Yekutieli)). (D) Quantification of the HSC population in cluster 17 of the control vs. GATA Pl1-KO placentae. The total number of cells in each sample type were used to calculate the percentage ((Mean±s.e.m., n=2 for each sample type). (E) Endothelial marker *Kdr* is highly expressed in cluster 15, unlike cluster 17, indicating hematoendothelial nature of the cells belonging to cluster 15. This cluster also harbors cells positive for Arterial-specific marker *Dll4* and placental hemogenic endothelium marker *Prom1*. (F) Flow cytometry analyses of Lin^-^ CD45^-^ CD34^+^ KDR^+^ cells show significant increase in the absence of pTGC-sepcific GATA factors. (G) Box plot of the flow analysis in the panel F. The percentages were calculated against total number of viable cells per sample (Mean±s.em., n=19 for the control and n=18 for the GATA Pl1-KO, **P≤0.01, analyzed by two-tailed Student’s t- test).

A hallmark of fetal hematopoiesis is the interrelated development of vascular and hematopoietic systems where hemogenic endothelium and hemangioblasts serve as the precursors to the hematopoietic stem cell populations ^41^. Multiple studies have shown that the embryonic hematopoiesis is intricately connected to the vascular development where hemogenic endothelium, a part of the vascular endothelial cells, gives rise to the definitive hematopoietic precursors during mammalian development ^42–44^. Thus, the genetic signature of the cells in cluster 15, which is unique to the knockout samples, indicates a hematoendothelial cell population that still retains the HSC lineage markers. This cell population was also found to express hematoendothelial markers *Cdh5*, *Icam2*, *Cd40*, confirming the bipotent nature of these cells ^45–47^ (**Fig. S8**). As a high expression of the arterial-specific marker *Delta-like, 4* (*Dll4*) is essential for the segregation of the endothelial lineage from the hematopoietic lineage ^48,49^, we looked at the *Dll4* expression in this subset, and found that this cluster expresses a high level of *Dll4* (**Fig. 3E**). Curiously, cells in cluster 15 also showed expression of a recently identified placental hemogenic endothelium marker *Prom1* ^50,51^, further confirming the hematoendothelial nature of cluster 15 (**Fig. 3E-F**).

We further looked at the hematoendothelial population by further subjecting the placental cell suspension to flow analysis. We screened single-cell suspension from control and GATA-Pl1 KO placentae and filtered them against other hematopoietic lineage cells using antibody specific for CD45, a pan-hematopoietic surface marker that appears on the more mature HSCs ^52^, and a cocktail of hematopoietic lineage marker antibodies (Lin). These cells were further screened for cells expressing CD34 and KDR simultaneously (**Fig. S9**). The percentage of this hematoendothelial Lin^-^ CD45^-^ CD34^+^ KDR^+^ population showed significant increase in the GATA- Pl1 KO sample than the controls (**Fig. 3F, G**).

The result from the flow analysis mirrors the data from the scRNA-seq experiments (**Fig. 3C**) and further confirms that the loss of GATA factors in the parietal TGCs skews the hematopoietic-endothelial lineage specification in a developing placenta and leads to the arrest of hematoendothelial progenitor population compared to the control.

In the next step, we used the three surface markers CD34, KIT (c-KIT), and Ly6A (SCA-1), which together define the placental HSCs, as the determinant for the long-term reconstituting (LTR) HSC population. Again, the cells were selected for CD45^-^ and Lin^-^ markers. These cells were further subjected to flow analysis and were selected for KIT^+^ CD34^+^ Ly6A/E^+^ cells. In the GATA-Pl1 KO placental samples, the Lin^-^ CD45^-^ KIT^+^ CD34^+^ Ly6A/E^+^ cell population was significantly reduced compared to the control (**Fig. 4A**). In comparison to the total number of cells, the percentage of the HSC population in the KO placenta was significantly lower than that in the control population (**Fig. 4B**) which resembles the results from the scRNA-seq analyses (**Fig. 3D**).

**Fig. 4:**
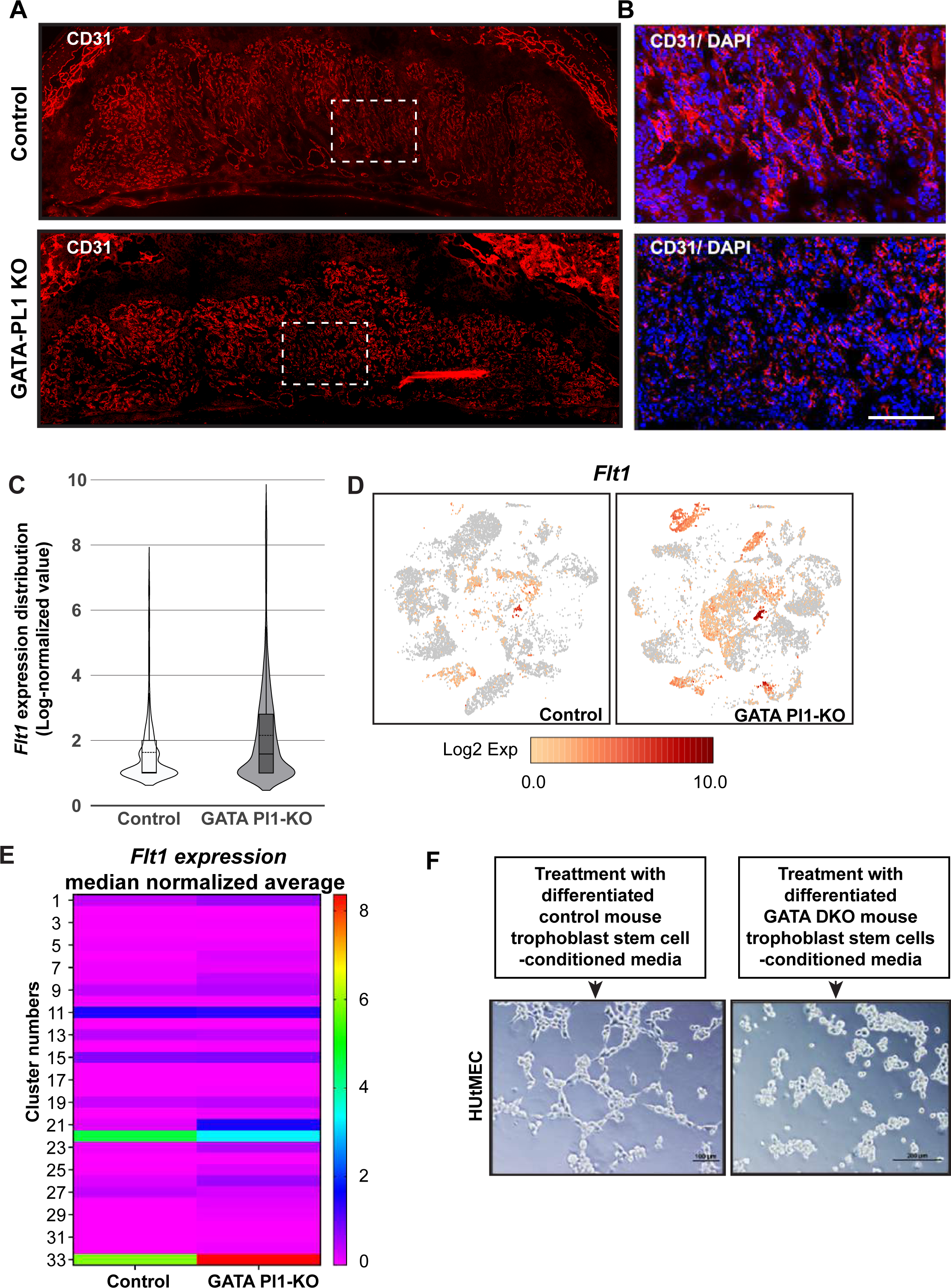
GATA factor loss in the TGCs results in the loss of the hematopoietic progenitor population. (A) Flow analysis data shows a significant reduction in the HSC population in the knockout placenta. E12.5 placental samples from GATA Pl1-KO and control littermate placentae were subjected to flow analysis. Samples were gated for viable cells, followed by gating for single cells. This was followed by selecting cells negative for lineage markers (Lin^-^) and subjecting them to CD45 negative selection. The Lin^-^ CD45^-^ population were further gated for CD34^+^ and Ly6A/E^+^ fraction. Finally these Lin^-^ CD45^-^ CD34^+^ Ly6A/E^+^ cells were gated for KIT^+^ population. The sample gating schematics are depicted using cell numbers representative of the respective panels. Black arrows show the flow of the steps. (B) Quantitation of the Lin^-^ CD45^-^ CD34^+^ KIT^+^ Ly6A/E^+^ population reveals significant loss of the HSC population in the KO sample. The counts are normalized against the number of viable cells and are presented as percentages (Mean±s.e.m., n=26 for the control and n=24 for the KO, **P≤0.01, analyzed by two-tailed Student’s t-test). (C) Micrographs show the formation of hematopoietic cell colonies on a methylcellulose plate. Colonies included were of CFU-G/M, BFU-E, and CFU-GEMM in nature. (D) Quantitation of the number of colonies in the control vs. GATA Pl1-KO samples. The counts are normalized against the number of cells plated for each sample (10,000 cells) (Mean±s.e.m., n=7 for each sample type, *P<0.05, analyzed by two-tailed Student’s t-test).

Next, we evaluated the differentiation potential of the GATA-Pl1 KO placental HSCs by using the colony-forming assay. Placental single-cell suspension from the control and GATA-Pl1 KO placentae were plated on methylcellulose medium containing a cocktail of Stem Cell Factor (SCF), IL-3, IL-6, and Erythropoietin. The placental samples gave rise to mostly granulocyte and mixed lineage erythroid, macrophage colonies. We observed a significant reduction in the number of colonies generated from the GATA-Pl1 KO placentae compared to the control (**Fig. 4C, D**).

It is speculated that the placental HSCs migrate and seed fetal liver along with the HSCs (Lin^-^ Ly6A/E^+^ KIT^+^ (LSK)) from the aorta-gonad-mesonephros (AGM) ^1,26,53,54^. In order to analyze the fetal liver HSC population, we performed FACS analysis of the E13.5 fetal liver of the GATA Pl1-KO embryos and compared them to the control. We found significant depletion of the LSK population in the GATA Pl1-KO fetal livers compared to the controls (**Fig. S10**), which validated our observation that the GATA Pl1-KO embryos show blood loss in their fetal liver (**Fig. 1G**).

Collectively these data prove that the GATA factor KO in the trophoblast giant cells negatively affects hematopoiesis in the placenta and fetal liver and results in the apparent blood loss phenotype. The loss of the HSC population was also accompanied by the incomplete segregation of the hematopoietic and endothelial lineage, indicating a loss of signaling network that balances and fine-tunes the hematopoietic versus endothelial cell lineage development in the placenta.

### TGC-specific loss of GATA factors affects placental vasculature development

Embryonic hematopoiesis and angiogenesis are tightly linked. Hemogenic endothelium in the vascular labyrinth gives rise to both the endothelial cell population as well as the hematopoietic cell population. As our study revealed a defective hematopoietic and endothelial lineage segregation due to TGC-specific GATA factor loss, we used our GATA-Pl1 KO placentae model to test placental angiogenesis. Anti-CD31 (PECAM-1) antibody, which marks early and mature endothelial cells, was used to stain the vasculature in the placental sections at E13.5. Compared to the control samples, the KO placentae showed gross disruption of the blood vessel architecture in the labyrinth (**Fig. 5A, B)**

**Fig. 5:**
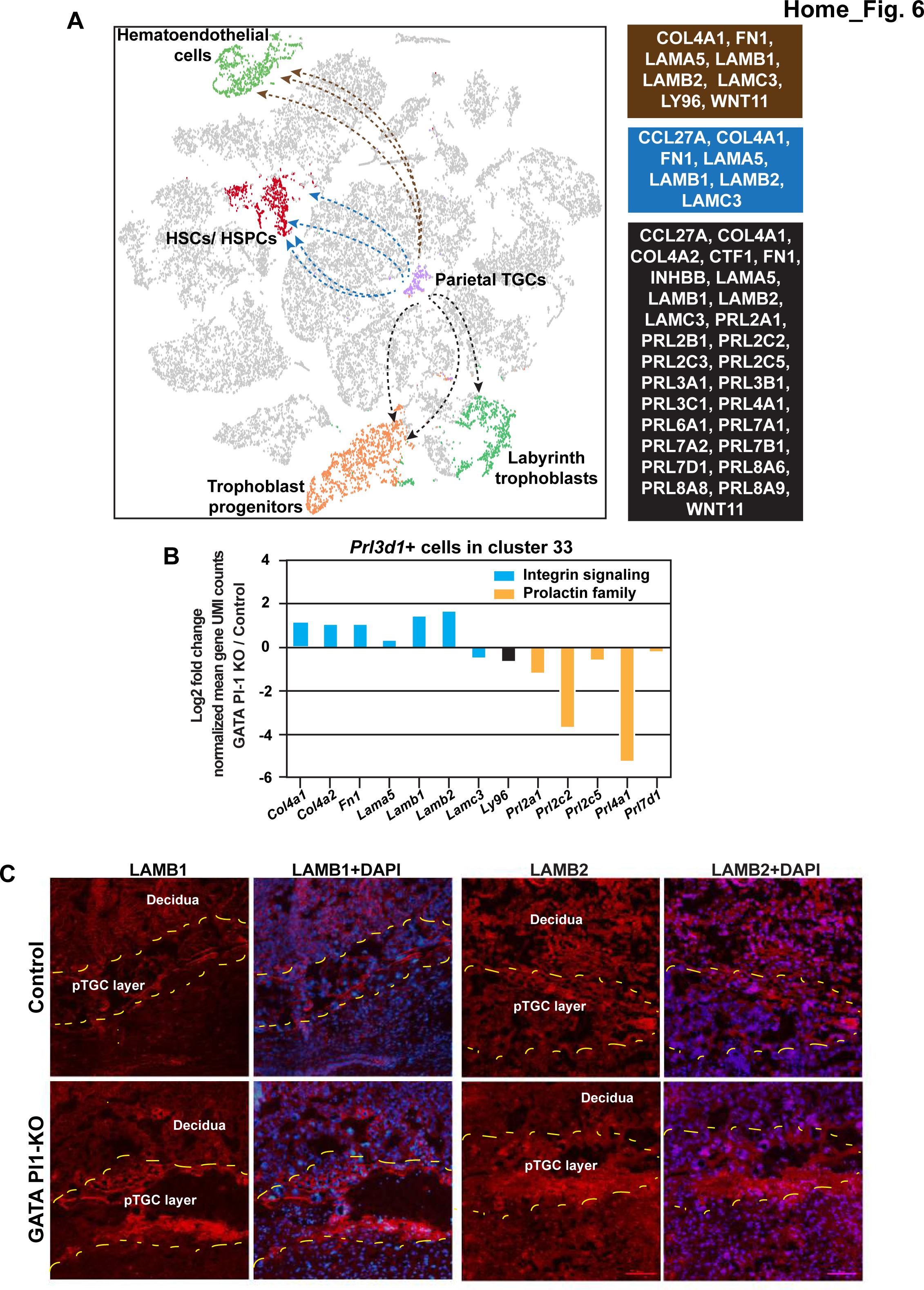
GATA factor-functions in the trophoblast giant cells are essential for the vasculature development at the maternal-fetal interface. (A-B) Immunostaining of the control and GATA Pl1-KO placenta with the endothelial cell marker CD31 shows abnormal vasculature development in the GATA Pl1-KO in the TGC layer (Scale Bar 200μm). (C) Violin plot shows *Flt1* expression increases in the GATA Pl1-KO placentae compared to the control. The plot (*log2* expression) and the values are derived from the scRNA-seq data and calculated using Loupe Browser (v 4.1.0) (10x Genomics). For the control, mean=1.63, median=1; for the KO mean=2.16, median=1.58. n=2 for both sample tytpes. (D) t-SNE plots for *Flt1* expression across the clusters show marked increase in the number of *Flt1* expressing cells in the GATA Pl1-KO placentae than control. Some of the major source of *Flt1* expression appears to be the TGC cluster 33, spiral artery associated TGC cluster 26, hematoendothelial cluster 15 and spongiotrophoblast trophoblast clsuters. (E) Heatmap of cluster-specific *Flt1* expression shows that the largest change in the *Flt1* expression occurs in the TGC cluster 33 (n=2 for each sample type). (F) In the presence of conditioned medium from GATA2/GATA3 double knockout (GATA DKO) trophoblast stem cells, HUtMECs fail to form vascular ring-like structures in a matrigel based vascular tube formation assays unlike when treated with conditioned medium from control trophoblast stem cells (n=3, Scale Bar 200μm).

Embryonic angiogenesis relies on a delicate balance of several key pro-angiogenic and anti- angiogenic factors ^55,56^. Among them anti-angiogenic factor soluble Vascular Endothelial Growth Factor (VEGF) receptor FLT1 (sFLT1) has been shown to play essential roles in the placental vascularization and angiogenesis ^57,58^. Our results showed upregulation of *Flt1* in the GATA-Pl1 KO placentae compared to the control (**Fig. 5C**). The sc-RNAseq data also showed that the highest upregulation of *Flt1* in the GATA Pl1-KO placentae was associated with cluster 33, previously identified in this study as the TGC cluster (**Fig. 5D, E**). Other prominent sources of Flt1 expression were the hematoendothelial cells (cluster 15), spongiotrophoblast cells in clusters 1, 9, 11, and spiral artery associated TGCs in cluster 26 (**Fig. 5D**).

To functionally test the increase in the *Flt1* level, we performed matrigel based vascular tube formation assays using Human Uterine Microvascular Endothelial Cells (HUtMEC). As differentiated trophoblast stem cells express antiangiogenic sFLT1, we harvested conditioned media from the differentiated *Gata2/Gata3* double knockout (GATA DKO) trophoblast stem cells and differentiated control trophoblast stem cells described in our earlier study ^19^. These conditioned media were added to the assays individually. Interestingly, HUtMECs in the presence of the conditioned media from the control trophoblast stem cells readily formed tubular structure while they failed to do so in the presence of conditioned media from GATA DKO trophoblast stem cells (**Fig. 5F**).

These findings indicate that the loss of GATA factors in the TGC layer results in the disruption of the delicate balance between the secreted angiogenic and anti-angiogenic factors by direct upregulation of *Flt1* expression in the TGC cells and by increasing the numbers of *Flt1* expressing trophoblast cells. This overall rise in the FLT1 level, in turn prevents proper development of the labyrinth vasculature in the placenta.

### GATA loss alters trophoblast giant cell-mediated paracrine signaling

TGCs are known to be a major source of autocrine and paracrine factors in the placenta ^32,59^. These factors, in turn, influence the development of the placenta as well as regulate numerous placental functions. In order to analyze the TGC-mediated signaling mechanism that regulates the development of placental trophoblast subpopulations and dictate hematoendothelial differentiation, we employed a recently reported bioinformatic tool PyMINEr ^60^. PyMINEr identifies receptor-receptor and ligand-receptor pairs by filtering out cell type-enriched receptors or secreted ligand genes. It then builds up a network of protein level interactions within and across all cell types by cross-referencing gene-gene pairs for physical protein-protein interactions. Finally, PyMINEr uses pathway analyses to identify the autocrine or paracrine signaling mechanism for these cell types. We used all the 33 clusters as input to PyMINEr to ensure correlation with the rest of our analyses. We identified several cross-cluster ligand- receptor pairs across our identified clusters. As we were interested in the paracrine signaling emanating from *Prl3d1*+ cluster 33, we evaluated all extracellular ligand pairs for corresponding receptor matches in other clusters. PyMINER analysis indicated the HSCs and HSPCs (cluster 17), and the hematoendothelial population (cluster 15), among the major paracrine signaling targets for extracellular factors expressed by the *Prl3d1*+ cluster 33 (**Fig. 6A**). This analysis also suggested laminin subunits *Lamb1*, *Lamb2*, *Lamc3*, *Lama5*, Collagen type IV alpha 1 (*Col4a1*), Fibronectin 1 (*Fn1*), and *Ccl27a* as possible major paracrine regulators acting on cluster 17 (**Dataset 4**). Similarly, our data suggested *Lamb1*, *Lamb2*, *Lamc3*, *Lama5*, Collagen type IV alpha 1 (*Col4a1*), Fibronectin 1 (*Fn1*), Lymphocyte antigen 96 (*Ly96*) and *Wnt11* to be the major potential paracrine factors acting on the hematoendothelial cluster 15 (**Fig. 6A**). When these genes were subjected to PANTHER pathway analysis, we observed significant enrichment of the Integrin signaling pathway in both the cases (**Fig. S11A**). The integrin signaling pathway has been implicated in both mouse and human placental development ^61^.

**Fig. 6:**
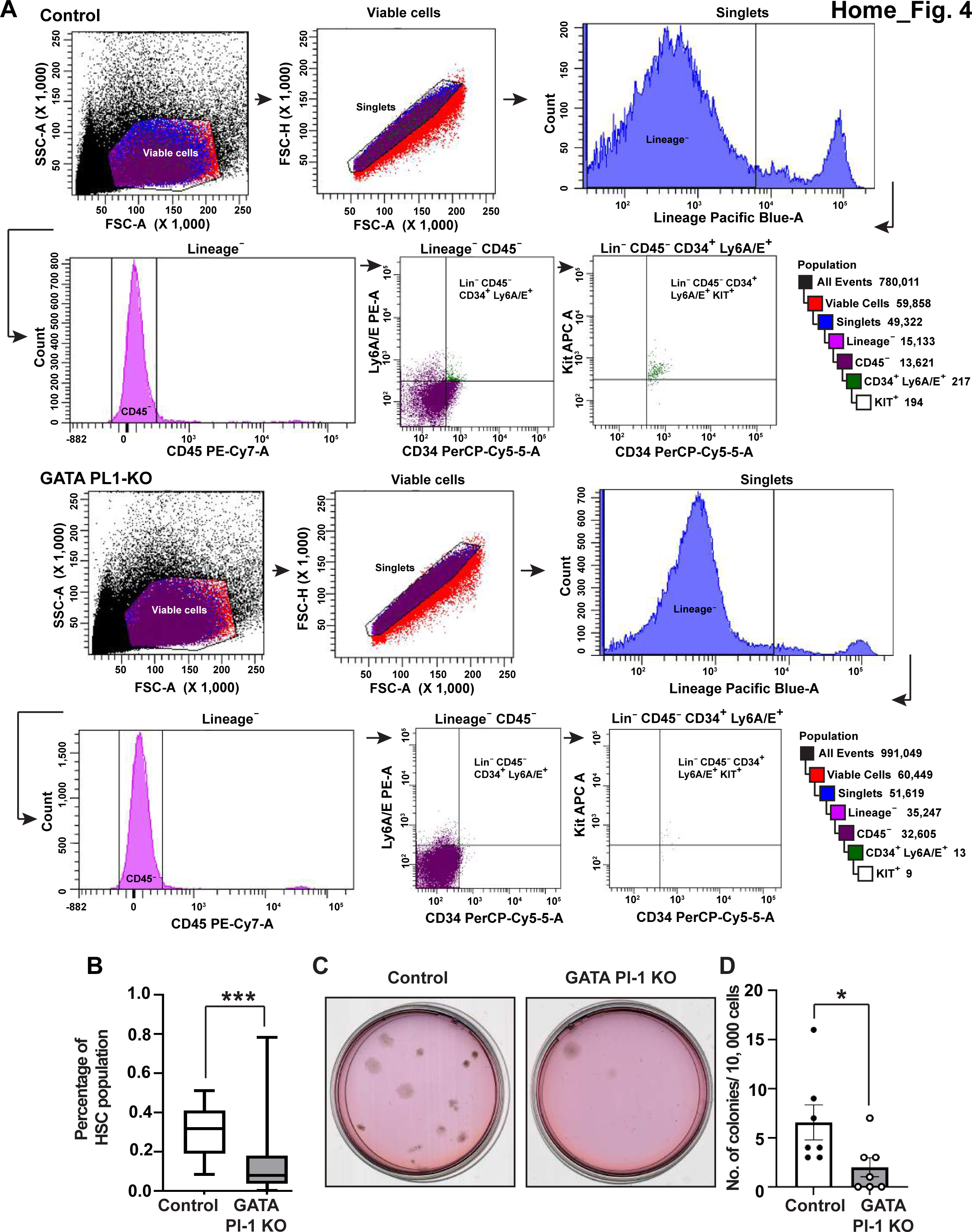
T**h**e **GATA factors regulate parietal TGC-specific paracrine factors and help maintain hematopoietic-angiogenic balance.** (A) PyMINEr based prediction of the enriched paracrine ligands expressed by cells in cluster 33 shows receptor matched targets in four major clusters that include HSPC/ HSC containing cluster 33, hematoendothelial cluster 15, trophoblast progenitor cluster 6 and labyrinthine trophoblast cluster 16. Major signaling pathways involved Integrin signaling pathway members lamin and collagen group 4 of proteins and prolactin family of hormones. (B) *Prl3d1* expressing cells in cluster 33 were further analyzed for the level of paracrine factors expression. While almost all the Integrin pathway members were upregulated, all the prolactin family members showed downregulation in the GATA Pl1-KO samples compared to the control. Average log2 fold change expression values are plotted. (C) Immunostaining of E13.5 placental sections with LAMBI and LAMB2 antibodies show increased expression for both the proteins in the pTGCs of the KO samples than controls (Scale Bar 100μm).

Interestingly, it has been shown that knocking out Integrin Subunit Alpha 5 (*Itga5*) in mice results in embryonic lethality for a large number of the embryos and shows poor development of the placental labyrinth and poor interdigitation of fetal and maternal vessels ^62^. Also, knocking out Integrin Subunit Alpha 7 (*Itga7*) in mice results in defective placental structures, including infiltration of the spongiotrophoblast layer into the placental labyrinth ^63^.

Moreover, our data pointed to two trophoblast clusters, progenitor trophoblasts (cluster 6), and labyrinthine trophoblasts (cluster 16) as the potential targets for 28 paracrine factors expressed by cluster 33 (**Fig. 6A**). These groups of paracrine factors included the Integrin signaling pathway members identified above, along with 16 prolactin family of hormones and Inhibin beta B (*Inhbb*) (**Dataset 4**). While the Prolactin family of genes has been studied extensively in the context of placental development and function ^64,65^, *Inhbb* has been implicated in preeclampsia66.

We analyzed these genes for GATA factor occupancy using our previously published data ^19^. Interestingly, we found about half of these genes (*Col4a1*, *Col4a2*, *Fn1*, *Inhbb*, *Lama5*, *Lamb1*, *Lamb2*, *Lamc3*, *Ly96*, *Prl2a1*, *Prl7d1, Wnt11*) to be putative GATA targets (**Fig. S11B-C**). Further analysis with the *Prl3d1*-positive cells in the cluster 33 showed upregulation of *Col4a1*, *Col4a2*, *Fn1*, *Inhbb*, *Lama5*, *Lamb1*, *Lamb2*, *Lamc3*, while downregulation of *Prl2a1*, *Prl2c2*, *Prl2c5*, *Prl4a1*, *Prl7d1* (**Fig. 6B**).

Among the suggested paracrine factors associated with the pTGCs in the GATA Pl1-KO placentae, we tested the expression level of LAMB1 and LAMB2. Immunostaining revealed increased expression levels of these two factors in the KO samples compared to the controls (**Fig. 6C**).

Collectively, these results suggest that inhibition of the Integrin signaling pathway in the parietal TGCs might be critical for hematoendothelial differentiation and maintaining hematopoietic- angiogenic balance in the developing mouse placenta. Our data also indicated that repression of the integrin signaling pathway along with the activation of the prolactin signaling might play a crucial part in the trophoblast lineage differentiation and helps maintain the proper ratio of placental trophoblast subtypes (**Fig. 7**).

**Fig. 7:**
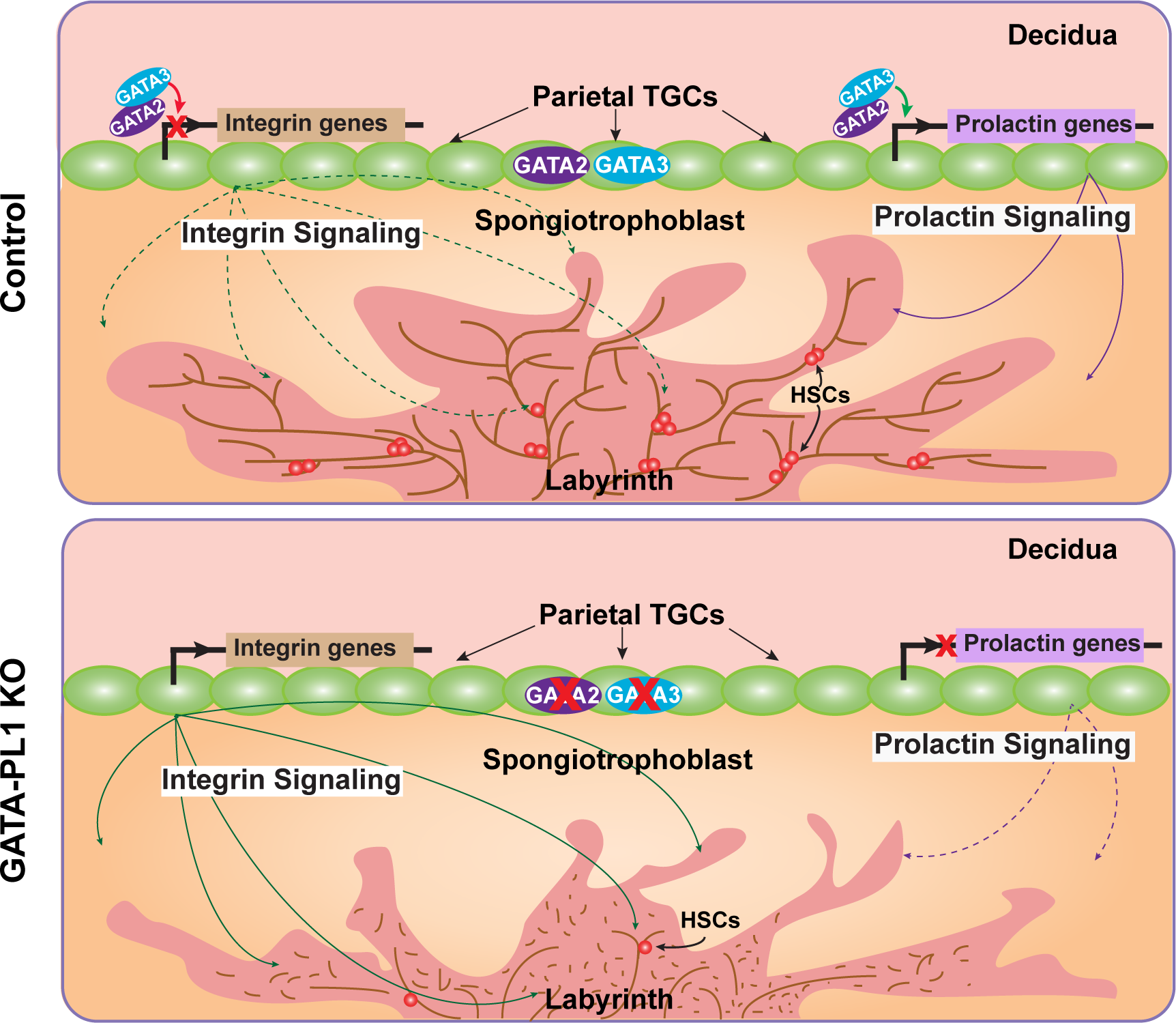
TGC-specific Integrin signaling and Prolactin signaling govern placental hematoendothelial differentiation and placental development. Schematic representation mid-gestation mouse placenta shows GATA factors regulate parietal-TGC-specific Integrin and Prolactin signaling. Repression of Integrin signaling and activation of Prolactin signaling by the GATA factors are achieved in the TGCs, which in turn helps angiogenic development, HSC development, and proper trophoblast differentiation in a control mouse placenta. In the absence of the GATA factors, an increase in the Integrin signaling and repression of the Prolactin signaling trigger loss of angiogenic branching and HSC population. This is accompanied by abnormal trophoblast differentiation and trophoblast subtype distribution.

## Discussion

GATA factor function in trophoblast differentiation and function has been subjected to numerous studies. Previously, we showed how GATA2 and GATA3 have overlapping functions during placental development, where the simultaneous loss of both the factors in all trophoblast cells leads to early embryonic death and placental defect with impaired hematopoiesis ^14,19^. It also revealed the loss of placenta layers, including the labyrinth region. Interestingly, the pan-trophoblast-specific knockout did not show significant change in the trophoblast giant cell numbers at the junctional zone.

As the placenta is a complex tissue consisting of diverse trophoblast subtypes as well as hematopoietic and endothelial cell populations, it is of utmost importance to categorize these cellular subtypes and define cell-cell interactions at the single-cell resolution. A recent study in mouse placenta employed single-nuclei RNAseq to highlight labyrinth development, but left out the role of the parietal TGCs ^67^. Although due to the inherent technical limtations of the 10X genomics sc-RNAseq platform we could only capture a small fraction of the syncytiotrophoblasts and TGCs, our analyses successfully identified the TGC cluster and the syncytial layers. Our sc-RNASeq data not only predicted the genetic signatures of the major trophoblast subtypes but also showed how the pTGC-specific transcriptional program governs the differentiation of the progenitor population and thereby altered the distribution of cells at different placental layers. Data from our *Gata2^f/f^*;*Gata3^f/f^; Pl1^Cre/wt^* mouse models showed that the loss of the GATA factors in the parietal TGC population severely reduced the placental size and also led to fetal growth restriction with significant phenotypic abnormality between E12.5 and E13.5. Importantly, we observed significant structural changes in the GATA-Pl1 KO placentae compared to the control. Molecular and morphological analyses revealed gross redistribution of different trophoblast progenitor cells, including *Prl2c2* expressing giant cells, *Tpbpa* expressing spongiotrophoblast progenitor, *Prl3b1* expressing secondary giant cell progenitors. While the number of *Epcam* expressing labyrinth progenitor cells showed a marked increase, we did not observe any significant enlargement of the labyrinth regions in the GATA-Pl1 KO placentae when normalized against the total placental area. In total, these findings reveal how different layers of trophoblast cells interact during placental development and influence the differentiation of trophoblast progenitors.

TGCs in the placenta secrete a lot of cytokines and hormones, which are not only crucial for trophoblast differentiation and placental development but are also critical for placental angiogenesis and hematopoiesis. Our study predicted possible mechanisms through which TGC-specific GATA factors dictate TGC-specific paracrine signaling and, in turn, regulate placental hematoendothelial niche. Curiously, the loss of TGC-specific GATA factors resulted in a unique cell population marked by the expression of *Kdr*, as well as *Kit*, *Cd34,* and *Ly6A*. The expression of both the endothelial gene and hematopoietic stem cell markers indicated towards the emergence of a hematoendothelial cell cluster. The complete absence of this cell population in the control placentae also indicated differentiation of these cells in the absence of TGC-specific GATA factors.

The GATA-Pl1 KO placentae were further characterized by the lack of proper angiogenic reoganization in the labyrinth. Our observation about the increased level of *Flt1* gives us a possible explanation about the angiogenic defect. The concomitant expansion of the *Tpbpa* expressing spongiotrophoblast in the GATA-Pl1 KO could also contribute to the impairment of the labyrinth development. We speculate that this loss in the surface area of exchange may result in the reduction of the nutrient supply and finally result in fetal growth restriction.

The integrin signaling pathway has been extensively studied in relation to the placental development as well as angiogenesis in both rodents and humans. Our novel finding suggests potential mechanisms by which inhibition of the integrin signaling pathway promotes placental hematoendothelial differentiation. This brings a new insight into how a small number of parietal trophoblast giant cells at the maternal-fetal interface can dictate the hematopoietic and angiogenic development in the placenta and thereby can affect fetal development.

Mouse embryogenesis involves vascularization on both the placental side as well as the decidual side. TGCs secrete a lot of angiogenic and anti-angiogenic paracrine factors, and in turn, regulate the decidual vascularization. However, how these factors dictate decidual vascularization, is poorly understood. It would be interesting to study how TGC-specific loss of GATA factors affect decidualization and associated vascular development. In this aspect, our TGC-specific GATA knockout mouse model holds great promise to help investigate the paracrine signaling from the TGC layers involved in the decidual angiogenesis.

GATA2 and GATA3 are evolutionarily conserved among mammals, and they are expressed in the human placenta. However, unlike the mouse placenta, human placental layers do not have trophoblast giant cells. In the absence of defined TGC subtypes, it is impossible to analyze the role of GATA factors in the placental development in humans. Moreover, ethical and logistical issues make it impossible to examine the role of GATA factors in human placental development *in vivo*. Nonetheless, the recent establishment of a human trophoblast cell line ^68^ has presented us with a new tool to examine the role of GATA factors in human placental development. As these cells can be readily differentiated *in vitro* to extravillous trophoblast and syncytiotrophoblast subtypes, they open the possibility of serving as models to study the role of GATA factors in the context of human placental development.

GATA2 and GATA3 are evolutionarily conserved among mammals, and they are expressed in the human placenta^19,69^. Although a human placenta does not have a Junctional zone, the TGC population resembles with the extravillous trophoblast cells (EVTs) of a human placenta. EVTs arise at the anchoring villi (which anchor the placenta with the uterine tissue) within a developing human placenta and invades into the uterine compartment^69^. A subset of EVT population migrates deeply into the uterus and upto the first third of the myometrium, where they differentiate to a giant cell population ^70,71^, which undergo endoreduplication, similar to the mouse parietal TGCs. These human invasive EVT-derived giant cells also mediate paracrine signaling to modulate the uterine microenviroment ^71^. Intriguingly both GATA2 and GATA3 are higly expressed in invasive EVTs at the human maternal-fetal interface. Thus, it will be interesting to find out how GATA2 and GATA3 regulate gene expression dynamics in human EVTs. Successful establishment of bona-fide human trophoblast stem cell lines ^68^ has presented us with a new tool to examine this aspect and opens up the possibility of serving as models to better understand the role of GATA factors in the context of human placental development.

## Materials and methods

### Generation of conditional knockout mice strains

All procedures were performed after obtaining IACUC approvals at the Univ. of Kansas Medical Center. Female *Gata2flox/flox* (*Gata2^f/f^*) mice ^72^ were mated with *Prl3d1tm1(cre)Gle* (*Pl1^Cre^*) male in order to generate *Gata2^f/+^; Pl1^Cre/wt^*. In the next step, *Gata2^f/+^; Pl1^Cre/wt^* female mice were bred with *Gata2^f/+^; Pl1^Cre/wt^* males to generate *Gata2^f/f^; Pl1^Cre/wt^*. Similarly female *Gata3flox/flox* (*Gata3^f/f^*) mice ^73^ were used to generate *Gata3^f/f^; Pl1^Cre/wt^*. In the next step, *Gata2^f/f^; Pl1^Cre/wt^* and Gata3^f/f^; *Pl1^Cre/wt^* mice were crossed to generate Gata2^f/+^;*Gata3^f/+^*; *Pl1^Cre/wt^*. Later *Gata2^f/+^;Gata3^f/+^; Pl1^Cre/wt^* males and females were crossed to generate *Gata2^f/f^;Gata3^f/f^; Pl1^Cre/wt^* strain. Further crosses with Gt(ROSA)26Sortm4(ACTB-tdTomato,-EGFP)Luo/J, (also known as mT/mG) mouse strain was used to establish *Gata2^f/f^;Gata3^f/f^;mT/mG* mouse line.

### Embryo harvest and tissue isolation

Animals were euthanized on desired day points, as indicated in the main text. Uterine horns and conceptuses were photographed. Conceptuses were dissected to isolate embryos, yolk sacs, and placentae. All embryos and placentae were photographed at equal magnification for comparison purposes.

Uteri containing placentation sites were dissected from pregnant female mice on E12.5 and E13.5 and frozen in dry ice-cooled heptane and stored at −80°C until used for histological analysis. Tissues were subsequently embedded in optimum cutting temperature (OCT) (Tissue-Tek) and were cryosectioned (10μm thick) for immunohistochemistry (IHC) studies using Leica CM-3050-S cryostat.

Placenta samples were carefully isolated, ensuring the decidual layer was peeled off. Individual samples were briefly digested in the presence of collagenase and were made into single-cell suspensions by passing them through a 40μm filter. These cell suspensions were further used for Flow analysis or FACS. Corresponding embryonic tissues were used to confirm genotypes. For scRNA-seq, these samples were further processed using Debris Removal Solution and Dead Cell Removal Kit (Miltenyi Biotec). Red blood cell depletion from the placental suspensions were carried out using anti-Mouse Ter-119 antibody (BD Biosciences) was used.

### Single Cell RNA-Sequencing and analysis

The transcriptomic profiles of mouse placental samples from two control (biological replicates) and two *Gata2/Gata3* double knockout (biological replicate) specimens were obtained using the 10x Genomics Chromium Single Cell Gene Expression Solution (10xgenomics.com). The primary analysis of the scRNA-seq data was performed using the 10x Genomics Cell Ranger pipeline (version 3.1.0). This pipeline performs sample de-multiplexing, barcode processing, and single-cell 3’ gene counting. The quality of the sequenced data was assessed using the FastQC software ^74^. Sequenced reads were mapped to the mouse reference genome (mm10) using the STAR software ^75^. Individual samples were aggregated using the “cellranger aggr” tool in Cell Ranger to produce a single feature-barcode matrix containing all the sample data. This process normalizes read counts from each sample, by subsampling, to have the same effective sequencing depth. The Cell Ranger software was used to perform two-dimensional PCA and t-SNE projections of cells, and k-means clustering. The 10x Genomics Loupe Cell Browser software (v 4.1.0) was used to find significant genes, cell types, and substructure within the single-cell data.

Digital Gene Expression matrices for each clusters identified using Cell Ranger, were individually fed into the scMCA pipeline (http://bis.zju.edu.cn/MCA/blast.html). scMCA output consisted of top markers for each clusters, predicted probabilities for the clusters alongwith their *p*-values. Clusters were identified using the smallest *p*-value. Where multiple equal probabilities existed, we calculated the abundance of the top markers in the respective clusters and also looked at the rest of the genes expressed in that cluster to attribute cell types in that cluster.

Using Ingenuity Pathway Analysis (Qiagen) software, we performed a core analysis of the significantly (*p* ≤ 0.05) upregulated transcripts in the *Prl3d1*+ cell clusters from the control and the GATA Pl1-KO scRNA-seq transcriptomes. The core analyses results were then compared and filtered for physiological functions related to hematopoiesis and angiogenesis to generate the heatmap.

PyMINEr analysis was performed using methods described by Tyler *et al.* ^60^. Clusters defined by the Cell Ranger were used as input. The receptor-ligand pairs were sorted on a PyMINER score. Pairs with a score more than 800 were retained to introduce high stringency. PANTHER pathway analysis was done using the predicted paracrine secreted ligands from cluster 33. Corresponding gene list were supplied as input.

### Cell culture and reagents

Mouse trophoblast stem cells (TSCs) were cultured with FGF4, Heparin, and MEF-conditioned medium (CM) according to protocol ^76^. *Gata2* and *Gata3* floxed alleles were efficiently excised from *Gata2^f/f^;Gata3^f/f^;UBC-cre/ERT2* (GATA DKO) TSCs by culturing the cells in absence of FGF4 and MEF-conditioned media and in the presence of tamoxifen (1 μg/ml) ^19^. Conditioned media was harvested upon removal of tamoxifen and was used for subsequent experiments.

### Flow analysis and sorting

For analyzing HSC population, placental single-cell suspensions were stained with APC-conjugated anti-mouse CD117 (c-Kit) (BioLegend), PerCP/Cy5.5-conjugated anti-mouse CD34 (BioLegend), PE-conjugated anti-mouse Ly-6A/E (Sca-1) (BioLegend), PE/Cy7 anti-mouse CD45 (BioLegend) and Pacific Blue-conjugated anti-mouse Lineage cocktail (BioLegend) monoclonal antibodies. Unstained, isotype and single-color controls were used for optimal gating strategy. Samples were run on either an LSRII flow cytometer or an LSRFortessa (BD Biosciences), and the data were analyzed using FACSDiva software.

Similarly, E13.5 fetal liver cells were also stained with APC-conjugated anti-mouse CD117 (c-Kit) (BioLegend), PE-conjugated anti-mouse Ly-6A/E (SCA-1) (BioLegend), and Pacific Blue-conjugated anti-mouse Lineage cocktail (BioLegend) monoclonal antibodies.

### Colony Formation Assay

Placenta cell suspensions were suspended in Iscove’s MDM with 2% fetal bovine serum and cultured using a MethoCult GF M3434 Optimum kit (STEMCELL Technologies). 10,000 cells from each sample were plated in 35-mm culture dishes (STEMCELL Technologies) and incubated at 37°C in a humidified, 5% CO2 environment for 14 days. Colonies were observed and counted using an inverted microscope.

### *In-Vitro* Endothelial Network Assembly Assay on Matrigel

Endothelial network assembly was assayed by the formation of capillary-like structures by HUtMECs on Matrigel (BD Biosciences). Matrigel was diluted 1:1 with a supplement-free M200 medium, poured in 12-well plates, and allowed to solidify at 37 °C. Sub confluent HUtMECs were harvested and preincubated for 1 hour in growth supplement-free M200 medium in microcentrifuge tubes. An equal volume of M200 medium containing FGF2/EGF was added. In addition, conditioned medium from differentiated *Gata2^f/f^*;*Gata3^f/f^;UBC^CreERT2^* trophoblast stem cells cultured in the presence (control) and absence of tamoxifen (GATA DKO) were added. The cells were plated on Matrigel (1.5 × 105 cells/well) and incubated at 37 °C and photographed at different time intervals.

### Quantitative RT-PCR

Total RNA from cells was extracted with the RNeasy Mini Kit (Qiagen). Samples isolated using FACS were further processed using the PicoPure RNA Isolation kit. cDNA samples were prepared and analyzed by qRT-PCR following procedures described earlier ^14^.

### Genotyping

Genomic DNA samples were prepared using tail tissues or embryonic tissues from the mice using the REDExtract-N-Amp Tissue PCR kit (Sigma-Aldrich). Genotyping was done using REDExtract-N-Amp PCR ReadyMix (Sigma-Aldrich) and respective primers. Respective primers are listed in the materials and methods section.

### Immunofluorescence

For immunostaining with mouse tissues, slides containing cryosections were dried, fixed with 4% PFA followed by permeabilization with 0.25% Triton X-100 and blocking with 10% fetal bovine serum and 0.1% Triton X-100 in PBS. Sections were incubated with primary antibodies overnight at 4 °C, washed in 0.1% Triton X-100 in PBS. After incubation (1:400, one hour, room temperature) with conjugated secondary antibodies, sections were washed, mounted using an anti-fade mounting medium (Thermo Fisher Scientific) containing DAPI and visualized using Nikon Eclipse 80i fluorescent microscope.

*mT/mG* positive embryos and cryosections were imaged directly under a Nikon Eclipse 80i fluorescent microscope.

### In situ hybridization

E13.5 whole conceptus cryosections were subjected to staining using RNAscope 2.5 HD Detection Kit (ACD Bio, Newark, CA). RNAscope probe for *Tpbpa* was used to detect the junctional zone of the mouse placenta, while hematoxylin was used as the counterstain.

### Statistical analyses

Independent data sets were analyzed using GraphPad Prism software. Two-tailed Student’s t-tests were performed for significabce and the data are presented as mean±s.e.m.

### Primer List

### Data availability

Processed data can be accessed from the NCBI Gene Expression Omnibus database (accession number GSE163286).

## Acknowledgment

This study is supported by various core facilities, including the Genomics Core, the Imaging and Histology Core facility, and the Bioinformatics Core of the University of Kansas Medical Center.

## Data availability

All the sequencing data are available and will be uploaded in a public database upon acceptance of this manuscript.

## Contribution

R.P.K. and S.P. conceived, designed and performed the initial study. P.H., RPK and A. Ghosh, performed the main experiments. P.H., A. Ghosh., R.P.K., and S.R. analyzed data. S.G., P.H. and R. K analyzed genomics data. P.H. wrote the manuscript. PH and RPK Revised the manuscript. S.P. edited the final manuscript.

## References

1. 1. Gekas, C., Rhodes, K.E., Van Handel, B., Chhabra, A., Ueno, M., and Mikkola, H.K. (2010). Hematopoietic stem cell development in the placenta. Int J Dev Biol 54, 1089–1098. 10.1387/ijdb.103070cg.

2. Ottersbach, K., and Dzierzak, E. (2005). The murine placenta contains hematopoietic stem cells within the vascular labyrinth region. Developmental Cell 8, 377–387. DOI 10.1016/j.devcel.2005.02.001.

3. Rhodes, K.E., Gekas, C., Wang, Y., Lux, C.T., Francis, C.S., Chan, D.N., Conway, S., Orkin, S.H., Yoder, M.C., and Mikkola, H.K. (2008). The emergence of hematopoietic stem cells is initiated in the placental vasculature in the absence of circulation. Cell Stem Cell 2, 252–263. 10.1016/j.stem.2008.01.001.

4. Duley, L. (2009). The global impact of pre-eclampsia and eclampsia. Seminars in perinatology 33, 130–137. 10.1053/j.semperi.2009.02.010.

5. 5. Lindheimer, M.D., Roberts, J.M., Cunningham, F.G., and Chesley, L.C. (2009). Chesley’s hypertensive disorders in pregnancy, 3rd Edition (Academic Press/Elsevier).

6. Iliodromiti, Z., Antonakopoulos, N., Sifakis, S., Tsikouras, P., Daniilidis, A., Dafopoulos, K., Botsis, D., and Vrachnis, N. (2012). Endocrine, paracrine, and autocrine placental mediators in labor. Hormones (Athens) 11, 397–409.

7. Nishikawa, S.I. (2001). A complex linkage in the developmental pathway of endothelial and hematopoietic cells. Curr Opin Cell Biol 13, 673–678.

8. Koushik, S.V., Wang, J., Rogers, R., Moskophidis, D., Lambert, N.A., Creazzo, T.L., and Conway, S.J. (2001). Targeted inactivation of the sodium-calcium exchanger (Ncx1) results in the lack of a heartbeat and abnormal myofibrillar organization. FASEB J 15, 1209–1211.

9. Hirashima, M., Lu, Y., Byers, L., and Rossant, J. (2003). Trophoblast expression of fms-like tyrosine kinase 1 is not required for the establishment of the maternal-fetal interface in the mouse placenta. Proceedings of the National Academy of Sciences of the United States of America 100, 15637–15642. 10.1073/pnas.2635424100.

10. Achen, M.G., Gad, J.M., Stacker, S.A., and Wilks, A.F. (1997). Placenta growth factor and vascular endothelial growth factor are co-expressed during early embryonic development. Growth Factors 15, 69–80. 10.3109/08977199709002113.

11. Tsai, F.Y., Keller, G., Kuo, F.C., Weiss, M., Chen, J., Rosenblatt, M., Alt, F.W., and Orkin, S.H. (1994). An early haematopoietic defect in mice lacking the transcription factor GATA-2. Nature 371, 221–226.

12. Pandolfi, P.P., Roth, M.E., Karis, A., Leonard, M.W., Dzierzak, E., Grosveld, F.G., Engel, J.D., and Lindenbaum, M.H. (1995). Targeted disruption of the GATA3 gene causes severe abnormalities in the nervous system and in fetal liver haematopoiesis. Nature genetics 11, 40–44. 10.1038/ng0995-40.

13. Ray, S., Dutta, D., Rumi, M.A.K., Kent, L.N., Soares, M.J., and Paul, S. (2009). Context-dependent function of regulatory elements and a switch in chromatin occupancy between GATA3 and GATA2 regulate Gata2 transcription during trophoblast differentiation. The Journal of biological chemistry 284, 4978–4988. 10.1074/jbc.M807329200.

14. Home, P., Ray, S., Dutta, D., Bronshteyn, I., Larson, M., and Paul, S. (2009). GATA3 is selectively expressed in the trophectoderm of peri-implantation embryo and directly regulates Cdx2 gene expression. J Biol Chem 284, 28729–28737. 10.1074/jbc.M109.016840.

15. Ghosh, A., Kumar, R., Kumar, R.P., Ray, S., Saha, A., Roy, N., Dasgupta, P., Marsh, C., and Paul, S. (2024). The GATA transcriptional program dictates cell fate equilibrium to establish the maternal-fetal exchange interface and fetal development. Proc Natl Acad Sci U S A 121, e2310502121. 10.1073/pnas.2310502121.

16. Ng, Y.K., George, K.M., Engel, J.D., and Linzer, D.I.H. (1994). Gata Factor Activity Is Required for the Trophoblast-Specific Transcriptional Regulation of the Mouse Placental- Lactogen-I Gene. Development 120, 3257–3266.

17. Ma, G.T., Roth, M.E., Groskopf, J.C., Tsai, F.Y., Orkin, S.H., Grosveld, F., Engel, J.D., and Linzer, D.I. (1997). GATA-2 and GATA-3 regulate trophoblast-specific gene expression in vivo. Development (Cambridge, England) 124, 907–914.

18. Jackson, D., Volpert, O.V., Bouck, N., and Linzer, D.I. (1994). Stimulation and inhibition of angiogenesis by placental proliferin and proliferin-related protein. Science 266, 1581–1584. 10.1126/science.7527157.

19. Home, P., Kumar, R.P., Ganguly, A., Saha, B., Milano-Foster, J., Bhattacharya, B., Ray, S., Gunewardena, S., Paul, A., Camper, S.A., et al. (2017). Genetic redundancy of GATA factors in the extraembryonic trophoblast lineage ensures the progression of preimplantation and postimplantation mammalian development. Development 144, 876–888. 10.1242/dev.145318.

20. Simmons, D.G., Fortier, A.L., and Cross, J.C. (2007). Diverse subtypes and developmental origins of trophoblast giant cells in the mouse placenta. Dev Biol 304, 567–578. 10.1016/j.ydbio.2007.01.009.

21. Bany, B.M., and Cross, J.C. (2006). Post-implantation mouse conceptuses produce paracrine signals that regulate the uterine endometrium undergoing decidualization. Dev Biol 294, 445–456. 10.1016/j.ydbio.2006.03.006.

22. Ouseph, M.M., Li, J., Chen, H.Z., Pecot, T., Wenzel, P., Thompson, J.C., Comstock, G., Chokshi, V., Byrne, M., Forde, B., et al. (2012). Atypical E2F repressors and activators coordinate placental development. Dev Cell 22, 849–862. 10.1016/j.devcel.2012.01.013.

23. Jiang, X., Wang, Y., Xiao, Z., Yan, L., Guo, S., Wang, Y., Wu, H., Zhao, X., Lu, X., and Wang, H. (2023). A differentiation roadmap of murine placentation at single-cell resolution. Cell Discovery 9, 30. 10.1038/s41421-022-00513-z.

24. Simmons, D.G., and Cross, J.C. (2005). Determinants of trophoblast lineage and cell subtype specification in the mouse placenta. Developmental biology 284, 12–24. 10.1016/j.ydbio.2005.05.010.

25. Muzumdar, M.D., Tasic, B., Miyamichi, K., Li, L., and Luo, L. (2007). A global double-fluorescent Cre reporter mouse. Genesis 45, 593–605. 10.1002/dvg.20335.

26. Mikkola, H.K., and Orkin, S.H. (2006). The journey of developing hematopoietic stem cells. Development 133, 3733–3744. 10.1242/dev.02568.

27. El-Hashash, A.H.K., Warburton, D., and Kimber, S.J. (2009). Genes and signals regulating murine trophoblast cell development. Mechanisms of development 127, 1–20. 10.1016/j.mod.2009.09.004.

28. Bany, B.M., and Cross, J.C. (2006). Post-implantation mouse conceptuses produce paracrine signals that regulate the uterine endometrium undergoing decidualization. Developmental Biology 294, 445–456. 10.1016/j.ydbio.2006.03.006.

29. Carney, E.W., Prideaux, V., Lye, S.J., and Rossant, J. (1993). Progressive expression of trophoblast-specific genes during formation of mouse trophoblast giant cells in vitro. Mol Reprod Dev 34, 357–368. 10.1002/mrd.1080340403.

30. Parisi, T., Beck, A.R., Rougier, N., McNeil, T., Lucian, L., Werb, Z., and Amati, B. (2003). Cyclins E1 and E2 are required for endoreplication in placental trophoblast giant cells. EMBO J 22, 4794–4803. 10.1093/emboj/cdg482.

31. Han, X., Wang, R., Zhou, Y., Fei, L., Sun, H., Lai, S., Saadatpour, A., Zhou, Z., Chen, H., Ye, F., et al. (2018). Mapping the Mouse Cell Atlas by Microwell-Seq. Cell 173, 1307. 10.1016/j.cell.2018.05.012.

32. Simmons, D.G., Fortier, A.L., and Cross, J.C. (2007). Diverse subtypes and developmental origins of trophoblast giant cells in the mouse placenta. Developmental biology 304, 567–578. 10.1016/j.ydbio.2007.01.009.

33. Natale, B.V., Schweitzer, C., Hughes, M., Globisch, M.A., Kotadia, R., Tremblay, E., Vu, P., Cross, J.C., and Natale, D.R.C. (2017). Sca-1 identifies a trophoblast population with multipotent potential in the mid-gestation mouse placenta. Sci Rep 7, 5575. 10.1038/s41598-017-06008-2.

34. Muntener, M., and Hsu, Y.C. (1977). Development of trophoblast and placenta of the mouse. A reinvestigation with regard to the in vitro culture of mouse trophoblast and placenta. Acta Anat (Basel) 98, 241–252.

35. Linzer, D.I., and Fisher, S.J. (1999). The placenta and the prolactin family of hormones: regulation of the physiology of pregnancy. Mol Endocrinol 13, 837–840. 10.1210/mend.13.6.0286.

36. Yotsumoto, S., Shimada, T., Cui, C.Y., Nakashima, H., Fujiwara, H., and Ko, M.S.H. (1998). Expression of adrenomedullin, a hypotensive peptide, in the trophoblast giant cells at the embryo implantation site in mouse. Developmental Biology 203, 264–275. DOI 10.1006/dbio.1998.9073.

37. WeilerGuettler, H., Aird, W.C., Rayburn, H., Husain, M., and Rosenberg, R.D. (1996). Developmentally regulated gene expression of thrombomodulin in postimplantation mouse embryos. Development 122, 2271–2281.

38. Lee, S.J., Talamantes, F., Wilder, E., Linzer, D.I.H., and Nathans, D. (1988). Trophoblastic Giant-Cells of the Mouse Placenta as the Site of Proliferin Synthesis. Endocrinology 122, 1761–1768. DOI 10.1210/endo-122-5-1761.

39. Cross, J.C., Hemberger, M., Lu, Y., Nozaki, T., Whiteley, K., Masutani, M., and Adamson, S.L. (2002). Trophoblast functions, angiogenesis and remodeling of the maternal vasculature in the placenta. Mol Cell Endocrinol 187, 207–212.

40. Gekas, C., Dieterlen-Lievre, F., Orkin, S.H., and Mikkola, H.K.A. (2005). The placenta is a niche for hematopoietic stem cells. Developmental Cell 8, 365–375. DOI 10.1016/j.devcel.2004.12.016.

41. Bollerot, K., Pouget, C., and Jaffredo, T. (2005). The embryonic origins of hematopoietic stem cells: a tale of hemangioblast and hemogenic endothelium. APMIS 113, 790–803. 10.1111/j.1600-0463.2005.apm_317.x.

42. Shalaby, F., Rossant, J., Yamaguchi, T.P., Gertsenstein, M., Wu, X.F., Breitman, M.L., and Schuh, A.C. (1995). Failure of blood-island formation and vasculogenesis in Flk-1- deficient mice. Nature 376, 62–66. 10.1038/376062a0.

43. Shalaby, F., Ho, J., Stanford, W.L., Fischer, K.D., Schuh, A.C., Schwartz, L., Bernstein, A., and Rossant, J. (1997). A requirement for Flk1 in primitive and definitive hematopoiesis and vasculogenesis. Cell 89, 981–990. Doi 10.1016/S0092-8674(00)80283-4.

44. Choi, K., Kennedy, M., Kazarov, A., Papadimitriou, J.C., and Keller, G. (1998). A common precursor for hematopoietic and endothelial cells. Development 125, 725–732.

45. Kim, I., Yilmaz, O.H., and Morrison, S.J. (2005). CD144 (VE-cadherin) is transiently expressed by fetal liver hematopoietic stem cells. Blood 106, 903–905. 10.1182/blood-2004-12-4960.

46. Nasrallah, R., Fast, E.M., Solaimani, P., Knezevic, K., Eliades, A., Patel, R., Thambyrajah, R., Unnikrishnan, A., Thoms, J., Beck, D., et al. (2016). Identification of novel regulators of developmental hematopoiesis using Endoglin regulatory elements as molecular probes. Blood 128, 1928–1939. 10.1182/blood-2016-02-697870.

47. Kubo, A., Chen, V., Kennedy, M., Zahradka, E., Daley, G.Q., and Keller, G. (2005). The homeobox gene HEX regulates proliferation and differentiation of hemangioblasts and endothelial cells during ES cell differentiation. Blood 105, 4590–4597. 10.1182/blood-2004-10-4137.

48. Park, M.A., Kumar, A., Jung, H.S., Uenishi, G., Moskvin, O.V., Thomson, J.A., and Slukvin, II (2018). Activation of the Arterial Program Drives Development of Definitive Hemogenic Endothelium with Lymphoid Potential. Cell Rep 23, 2467–2481. 10.1016/j.celrep.2018.04.092.

49. Marcelo, K.L., Goldie, L.C., and Hirschi, K.K. (2013). Regulation of endothelial cell differentiation and specification. Circ Res 112, 1272–1287. 10.1161/CIRCRESAHA.113.300506.

50. 50. Pereira, C.F., Chang, B., Gomes, A., Bernitz, J., Papatsenko, D., Niu, X., Swiers, G., Azzoni, E., de Bruijn, M.F., Schaniel, C., et al. (2016). Hematopoietic Reprogramming In Vitro Informs In Vivo Identification of Hemogenic Precursors to Definitive Hematopoietic Stem Cells. Dev Cell 36, 525–539. 10.1016/j.devcel.2016.02.011.

51. Azevedo Portilho, N., and Pelajo-Machado, M. (2018). Mechanism of hematopoiesis and vasculogenesis in mouse placenta. Placenta 69, 140–145. 10.1016/j.placenta.2018.04.007.

52. 52. North, T.E., de Bruijin, M.F.T.R., Stacy, T., Talebian, L., Lind, E., Robin, C., Binder, M., Dzierzak, E., and Speck, N.A. (2002). Runx1 expression marks long-term repopulating hematopoietic stem cells in the midgestation mouse embryo. Immunity 16, 661–672. Doi 10.1016/S1074-7613(02)00296-0.

53. 53. Lee, L.K., Ueno, M., Van Handel, B., and Mikkola, H.K.A. (2010). Placenta as a newly identified source of hematopoietic stem cells. Curr Opin Hematol 17, 313–318. 10.1097/MOH.0b013e328339f295.

54. Mahony, C.B., and Bertrand, J.Y. (2019). How HSCs Colonize and Expand in the Fetal Niche of the Vertebrate Embryo: An Evolutionary Perspective. Front Cell Dev Biol 7. UNSP 34 10.3389/fcell.2019.00034.

55. Breier, G. (2000). Angiogenesis in embryonic development--a review. Placenta 21 *Suppl A*, S11-15.

56. Drake, C.J., and Fleming, P.A. (2000). Vasculogenesis in the day 6.5 to 9.5 mouse embryo. Blood 95, 1671–1679.

57. 57. Carmeliet, P., Moons, L., Luttun, A., Vincenti, V., Compernolle, V., De Mol, M., Wu, Y., Bono, F., Devy, L., Beck, H., et al. (2001). Synergism between vascular endothelial growth factor and placental growth factor contributes to angiogenesis and plasma extravasation in pathological conditions. Nat Med 7, 575–583. 10.1038/87904.

58. Palmer, K.R., Tong, S., and Kaitu’u-Lino, T.J. (2017). Placental-specific sFLT-1: role in pre-eclamptic pathophysiology and its translational possibilities for clinical prediction and diagnosis. Mol Hum Reprod 23, 69–78. 10.1093/molehr/gaw077.

59. Hu, D., and Cross, J.C. (2010). Development and function of trophoblast giant cells in the rodent placenta. Int J Dev Biol 54, 341–354. 10.1387/ijdb.082768dh.

60. Tyler, S.R., Rotti, P.G., Sun, X., Yi, Y., Xie, W., Winter, M.C., Flamme-Wiese, M.J., Tucker, B.A., Mullins, R.F., Norris, A.W., and Engelhardt, J.F. (2019). PyMINEr Finds Gene and Autocrine-Paracrine Networks from Human Islet scRNA-Seq. Cell Rep 26, 1951–1964 e1958. 10.1016/j.celrep.2019.01.063.

61. Lee, C.Q.E., Turco, M.Y., Gardner, L., Simons, B.D., Hemberger, M., and Moffett, A. (2018). Integrin alpha2 marks a niche of trophoblast progenitor cells in first trimester human placenta. Development 145. 10.1242/dev.162305.

62. Bader, B.L., Rayburn, H., Crowley, D., and Hynes, R.O. (1998). Extensive vasculogenesis, angiogenesis, and organogenesis precede lethality in mice lacking all alpha v integrins. Cell 95, 507–519. 10.1016/s0092-8674(00)81618-9.

63. Welser, J.V., Lange, N.D., Flintoff-Dye, N., Burkin, H.R., and Burkin, D.J. (2007). Placental defects in alpha7 integrin null mice. Placenta 28, 1219–1228. 10.1016/j.placenta.2007.08.002.

64. Wiemers, D.O., Shao, L.J., Ain, R., Dai, G., and Soares, M.J. (2003). The mouse prolactin gene family locus. Endocrinology 144, 313–325. 10.1210/en.2002-220724.

65. Simmons, D.G., Rawn, S., Davies, A., Hughes, M., and Cross, J.C. (2008). Spatial and temporal expression of the 23 murine Prolactin/Placental Lactogen-related genes is not associated with their position in the locus. BMC Genomics 9, 352. 10.1186/1471-2164-9- 352.

66. Johnson, M.P., Brennecke, S.P., East, C.E., Goring, H.H., Kent, J.W., Jr., Dyer, T.D., Said, J.M., Roten, L.T., Iversen, A.C., Abraham, L.J., et al. (2012). Genome-wide association scan identifies a risk locus for preeclampsia on 2q14, near the inhibin, beta B gene. PLoS One 7, e33666. 10.1371/journal.pone.0033666.

67. Marsh, B., and Blelloch, R. (2020). Single nuclei RNA-seq of mouse placental labyrinth development. Elife 9. 10.7554/eLife.60266.

68. Okae, H., Toh, H., Sato, T., Hiura, H., Takahashi, S., Shirane, K., Kabayama, Y., Suyama, M., Sasaki, H., and Arima, T. (2018). Derivation of Human Trophoblast Stem Cells. Cell Stem Cell 22, 50–63 e56. 10.1016/j.stem.2017.11.004.

69. Paul, S., Home, P., Bhattacharya, B., and Ray, S. (2017). GATA factors: Master regulators of gene expression in trophoblast progenitors. Placenta 60 *Suppl 1*, S61–S66. 10.1016/j.placenta.2017.05.005.

70. Pollheimer, J., Vondra, S., Baltayeva, J., Beristain, A.G., and Knofler, M. (2018). Regulation of Placental Extravillous Trophoblasts by the Maternal Uterine Environment. Front Immunol 9, 2597. 10.3389/fimmu.2018.02597.

71. Arutyunyan, A., Roberts, K., Troule, K., Wong, F.C.K., Sheridan, M.A., Kats, I., Garcia-Alonso, L., Velten, B., Hoo, R., Ruiz-Morales, E.R., et al. (2023). Spatial multiomics map of trophoblast development in early pregnancy. Nature 616, 143–151. 10.1038/s41586-023-05869-0.

72. Charles, M.A., Saunders, T.L., Wood, W.M., Owens, K., Parlow, A.F., Camper, S.A., Ridgway, E.C., and Gordon, D.F. (2006). Pituitary-specific Gata2 knockout: effects on gonadotrope and thyrotrope function. Mol Endocrinol 20, 1366–1377. 10.1210/me.2005-0378.

73. Zhu, J., Min, B., Hu-Li, J., Watson, C.J., Grinberg, A., Wang, Q., Killeen, N., Urban, J.F., Guo, L., and Paul, W.E. (2004). Conditional deletion of Gata3 shows its essential function in T(H)1-T(H)2 responses. Nature Immunology 5, 1157–1165.

74. 74. Andrews, S. (2010). FastQC: a quality control tool for high throughput sequence data. Available online at: http://www.bioinformatics.babraham.ac.uk/projects/fastqc.

75. Dobin, A., Davis, C.A., Schlesinger, F., Drenkow, J., Zaleski, C., Jha, S., Batut, P., Chaisson, M., and Gingeras, T.R. (2013). STAR: ultrafast universal RNA-seq aligner. Bioinformatics 29, 15–21. 10.1093/bioinformatics/bts635.

76. Tanaka, S., Kunath, T., Hadjantonakis, A.K., Nagy, A., and Rossant, J. (1998). Promotion of trophoblast stem cell proliferation by FGF4. Science 282, 2072–2075.

