## Supporting Figures for "A Single Trophoblast Layer Acts as the Gatekeeper at the Endothelial-Hematopoietic Crossroad in the Placenta"

**Fig. S1**

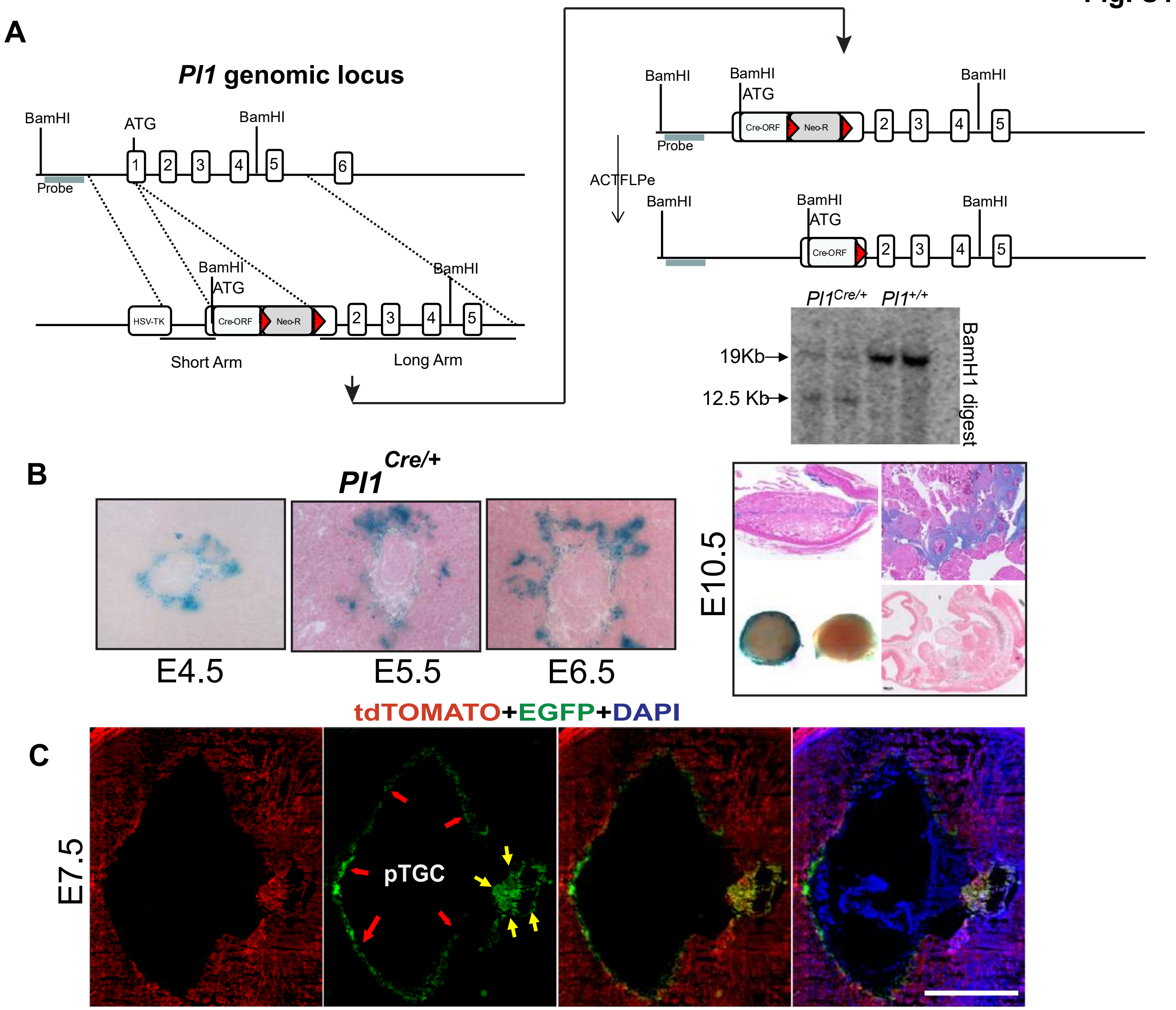


**Fig. S1.** (A) Schematics showing the plan to generate the *Pl1*$^{Cre/+}$ mouse line. Briefly, RPCI-22 (129S6/SvEvTac) Mouse BAC Library (BPRC, Children's Hospital Oakland Research Institute) was screened to identify BAC clone containing *Pl1* gene. Gene targeting construct was generated to include close to 4kb short arm and 8kb long arm using standard cloning and recombineering protocols. Frt sequence flanked Neomycin selection (NeoR) cassette was used to facilitate positive selection while thymidine kinase (TK) cassette was utilized for gancyclovir based negative selection. Linearized targeting vector was used for gene targeting using TC1 embryonic stem cells (Nationwide Children's Hospital Research Core Facility). Positive clones were identified by southern blot screening as shown in figure and were injected into C57BL/6 blastocyst for generating chimeras. Chimeric mice were bred with Black Swiss wild type mice to acquire germline transmission. These founders were further bred with Flpe transgenic mice (B6.Cg-Tg (ACTFLPe) FLPe: JAX Strain # 005703) to remove neomycin selection cassette and were back bred with Black Swiss wild type mice to remove Flpe transgene and to generate final experimental animals. (B) Temporal expression of *Pl1*$^{Cre}$ as visualized by LacZ reporter staining at E4.5, E5.5, E6.5, and E10.5 and (C) Cre recombinase activity allowing tdTOMATO to EGFP switch at E7.5 (red arrows pointing pTGCs and yellow arrows indicating developing placenta). Scale bar 1000um.

**Litter analysis**

*Gata2*$^{f/f}$;*Gata3*$^{f/f}$;*Pl1*$^{Cre/wt}$ **(male) X** *Gata2*$^{f/f}$;*Gata3*$^{f/f}$ **(female)**

**A**

| Gestational day | Number of litters analyzed | Total number of implantation sites | Cre-negative embryos | Cre-positive GATA Pl1-KO embryos | |
|---|---|---|---|---|---|
| | | | | Non-resorbed embryos | Extremely small, dead, resorbed embryos |
| **E12.5** | 14 | 126 | 67 (53.17%) | 40 (31.74%) | 19 (15.07%) |
| **E13.5** | 13 | 119 | 62 (52.10%) | 34 (28.57%) | 23 (19.32%) |

**B**

| Non-resorbed GATA Pl1-KO embryos (E12.5-E13.5) | | |
|---|---|---|
| Total | Embryos with no apparent phenotypes | Embryos with visible smaller size/ blood loss phenotypes |
| 40 | 11 (27.50%) | 29 (72.50%) |
| 34 | 7 (20.58%) | 27 (79.41%) |

**C**

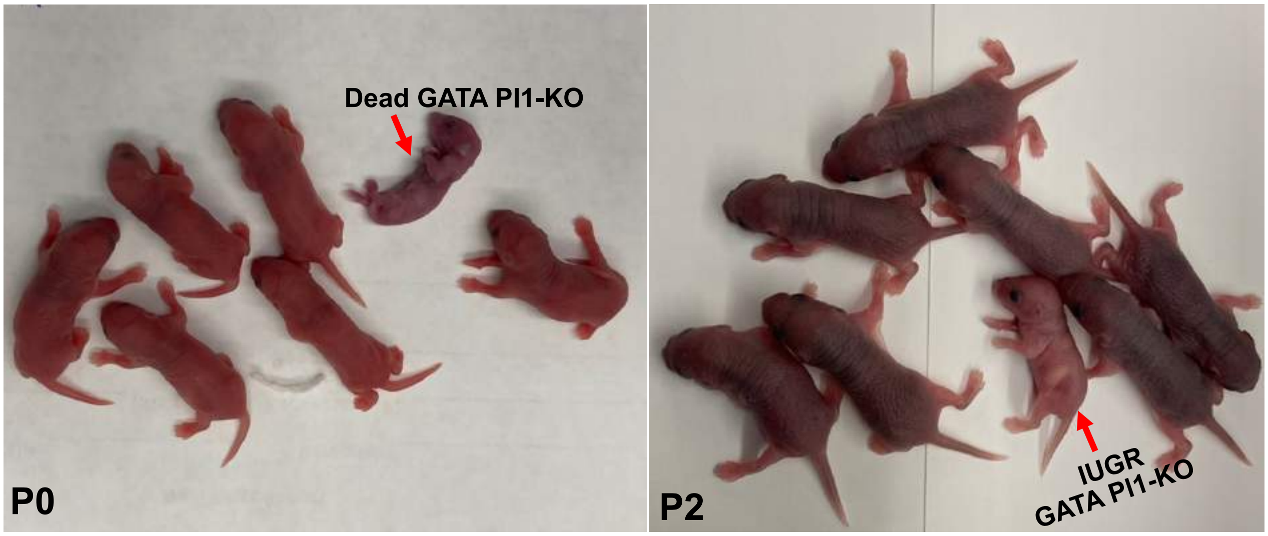


**Fig. S2.** Litter analyses from matings between *Gata2*$^{f/f}$;*Gata3*$^{f/f}$; *Pl1Cre* $^{+/wt}$ males and *Gata2*$^{f/f}$;*Gata3*$^{f/f}$ females. (A) 14 litters from gestational day E12.5 and 13 litters from gestational day E13.5 were analyzed in total. PCR were done using part of the embryo proper to confirm the genotyping. (B) Further distribution of the non-resorbed Cre-positive GATA Pl1-KO embryos. (C) Some of the GATA Pl1-KO pups were delivered dead (P0) and even some of the survived pups showed IUGR phenotype at postnatal day 2 (P2).

**Fig. S3**

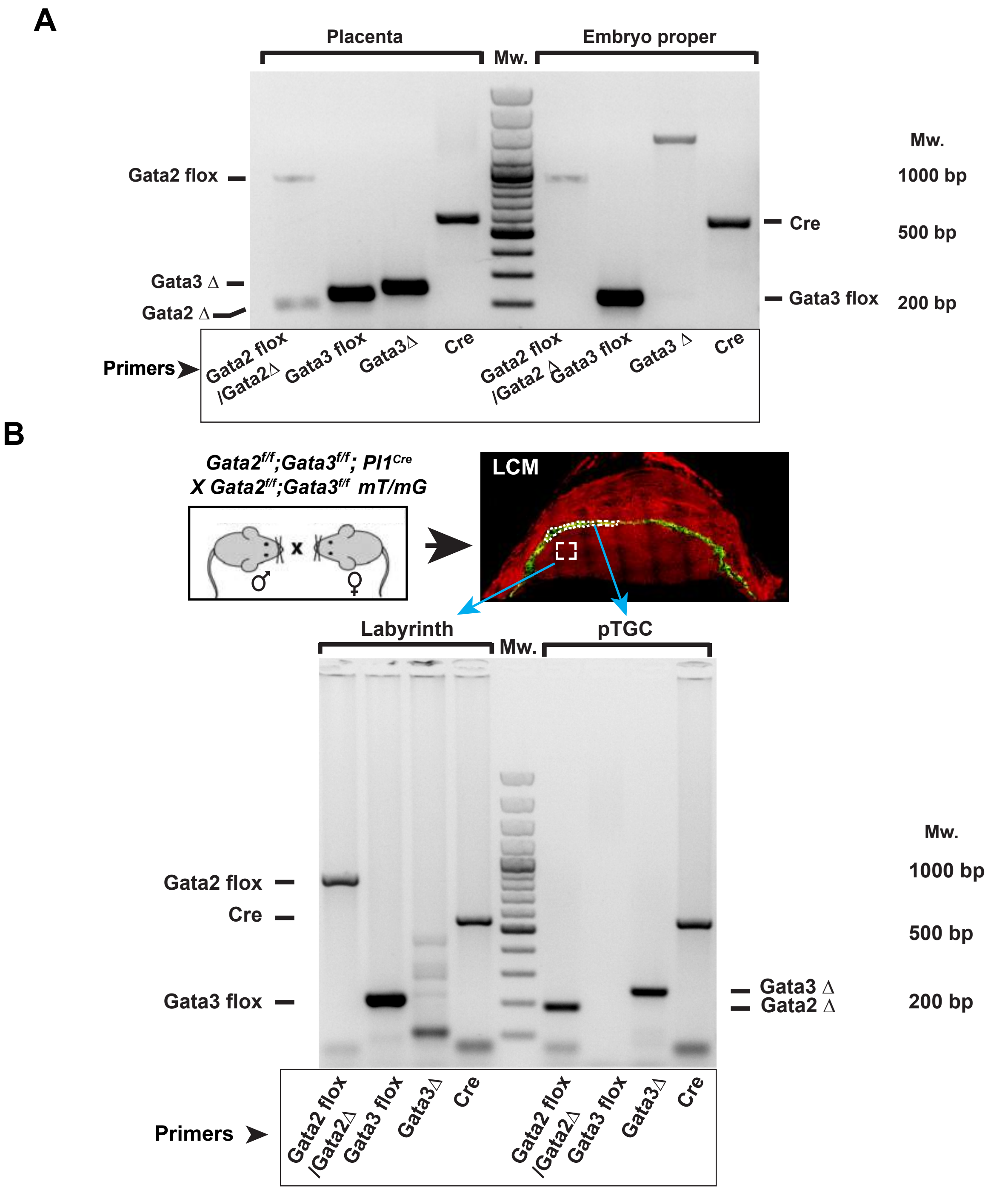


**Fig. S3.** (A) Genotyping results from the whole placental tissue and corresponding embryo proper of a *Gata2*$^{f/f}$;*Gata3*$^{f/f}$; *Pl1Cre* $^{+/wt}$ embryo confirms that the deletion of the GATA alleles were confined to the placenta. Only the placental samples showed deletion bands corresponding to *Gata2* and *Gata3* deletions. Presence of the *Gata2* $^{flox}$ band in the placental sample is the result of the contribution from other trophoblast and non-trophoblast cell types. (B) Placental sections from E12.5 *Gata2*$^{f/f}$;*Gata3*$^{f/f}$; *Pl1Cre* $^{+/wt}$; *mT/mG* were subjected to laser capture microdissection followed by PCR. Samples were excised from the GFP positive parietal TGCs (pTGCs) while the control samples were excised from labyrinth region. Care was taken to not include any red fluorescent region while excising for the GFP positive sections. Samples were then used to extract DNA using REDExtract-N-Amp Tissue PCR kit (Sigma-Aldrich) and subjected to PCR. Only the pTGC samples showed deletion bands corresponding to *Gata2* and *Gata3,* confirming the specific nature of PL1-Cre mediated deletions. Moreover, the lack of flox bands in the LCM samples also show the high efficiency of the Cre-mediated deletions.

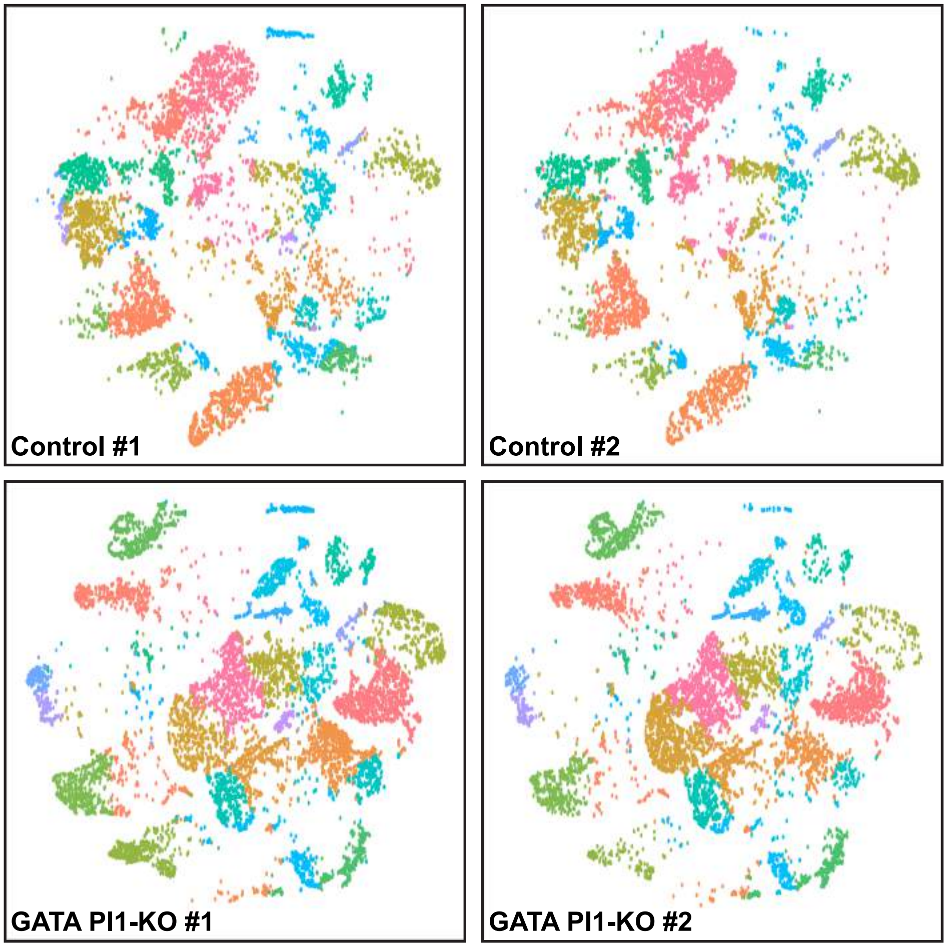


**Fig. S4.** t-SNE plot showing similar clustering patterns between two different control placental samples, and two different GATA PI1-KO placental samples in the scRNA-seq analyses.

**Fig. S5**

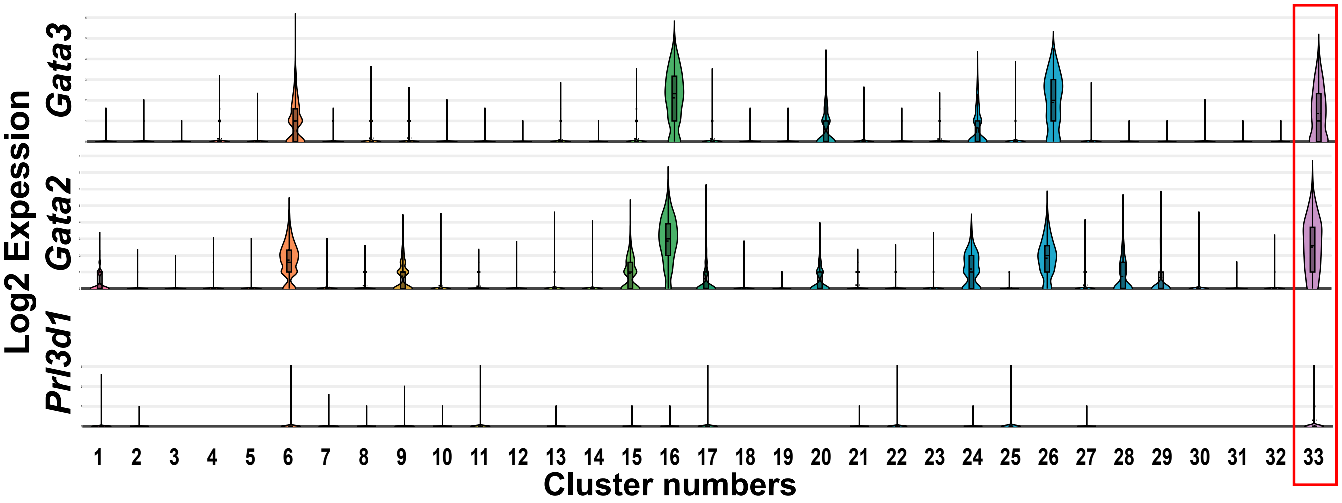


**Fig. S5.** Violin plots of expression of *Prl3d1*, *Gata2* and *Gata3* across all the clusters. Significant expression of *Gata2* and *Gata3* were observed predominantly in the cluster 33.

Fig. S6

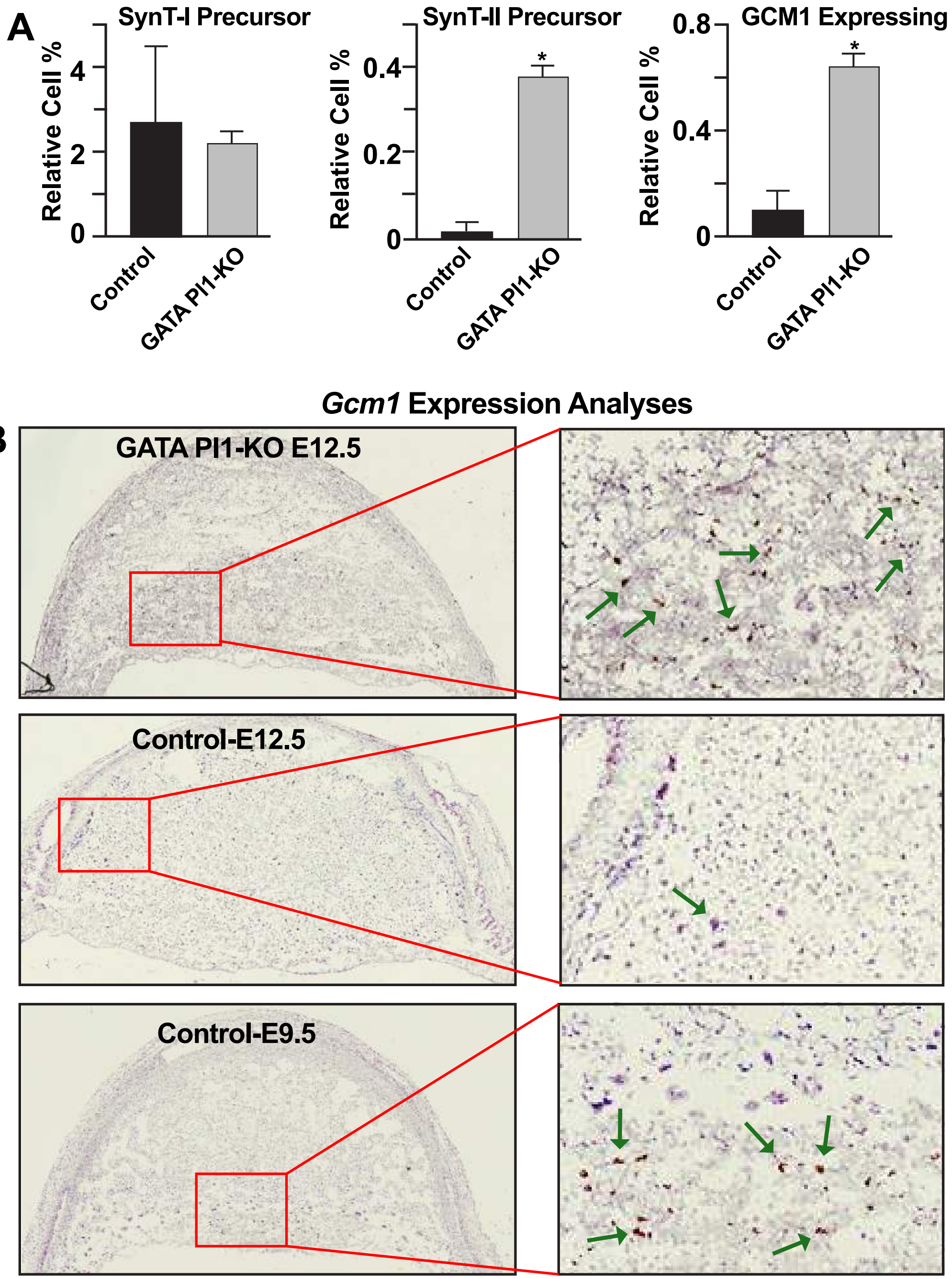


**Fig. S6. (A) Quantitation of relative cell percentage showing GATA2/3 transcription factors control SynT layer development and *Gcm1* expressing population and (B) Validation of *Gcm1* expressing trophoblast cells using RNAScope labeling of the *Gcm1* at different stages of placenta. Comparison of the relative abundance of Gcm1+ trophoblasts decline tremendously from e9.5 to e12.5, while *Gcm1* expression persists in the GATA-PI1 KO.**

**Fig. S7**

### IPA based Physiological Functions of *Prl3d1*+ cells

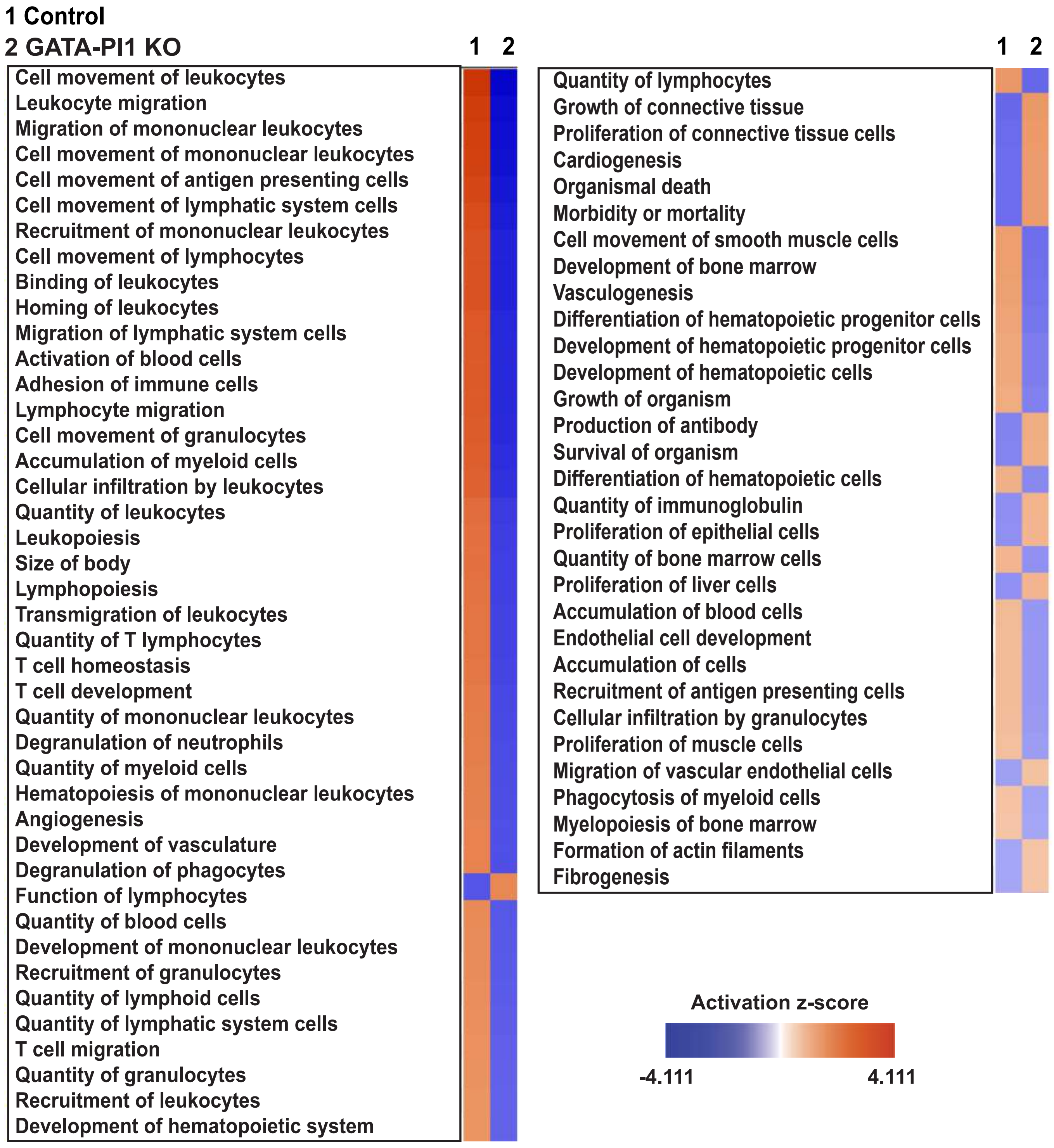


**Fig. S7.** Comparison of hematopoiesis and angiogenesis related physiological functions between the *Prl3d1* positive TGCs in control and GATA-Pl1 KO samples. Significantly upregulated gene expression (p<=0.05) of the *Prl3d1* positive cells from the control and the GATA-Pl1 KO samples (compared to the rest of the cells of the corresposnding placentate) were subjected to core analysis of the Ingenuity Pathway Analysis. Physiological functions related to hematopoiesis and angiogenesis were then compared to generate the heatmap.

Fig. S8

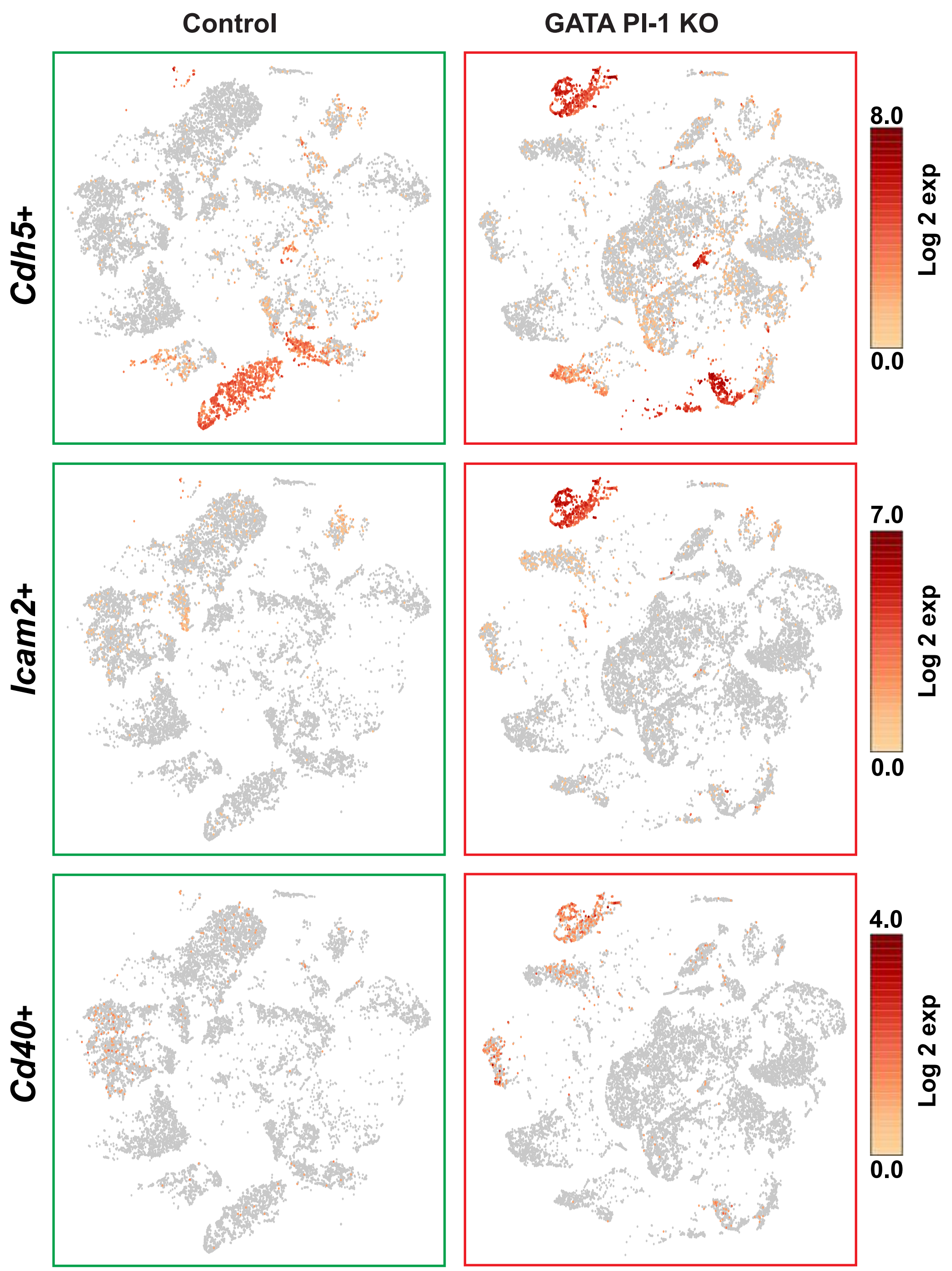


**Fig. S8.** Cluster 15, which is predominately populated in the GATA PI1-KO placentae shows high level of expression for hematoendothelial markers *Cdh5, Icam2* and *Cd40*.

**Fig. S9**

**Control**

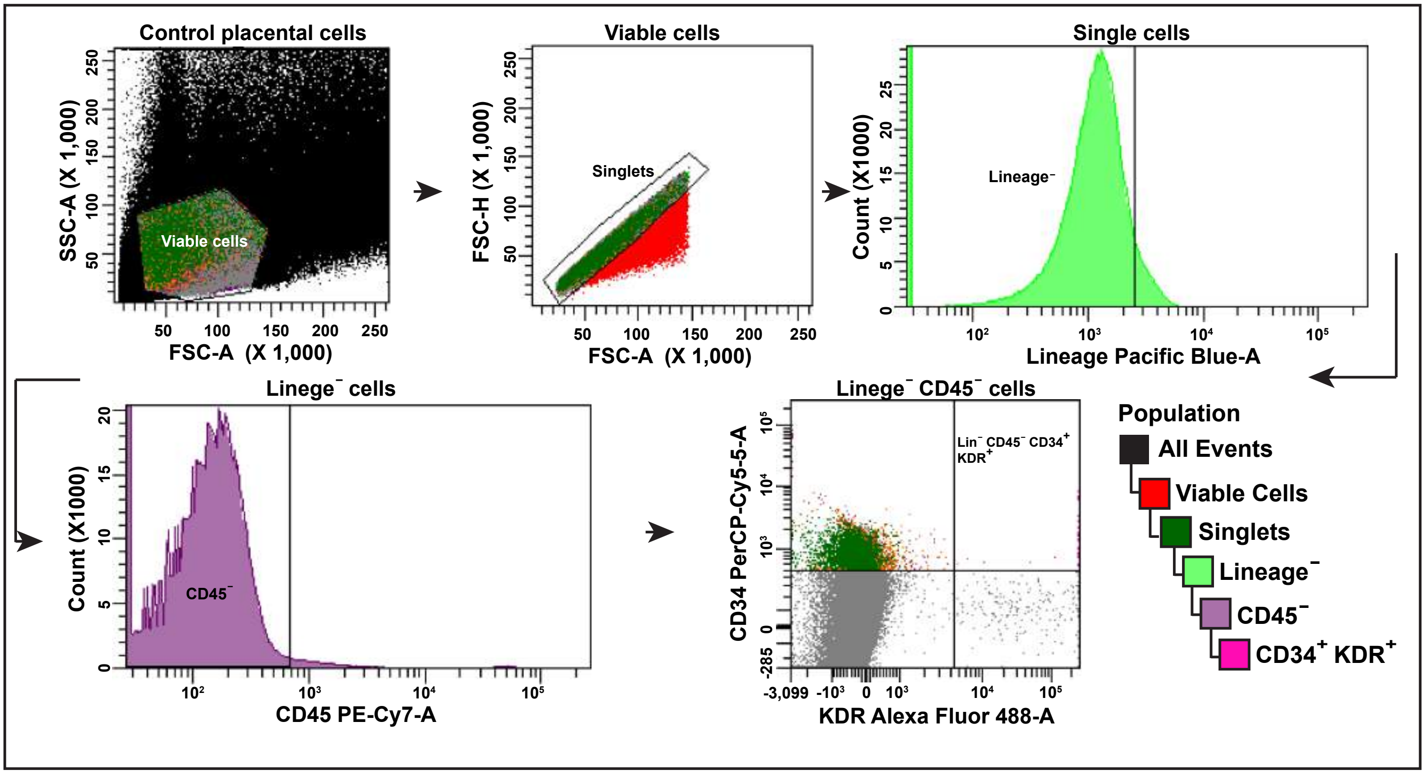


**GATA PL1-KO**

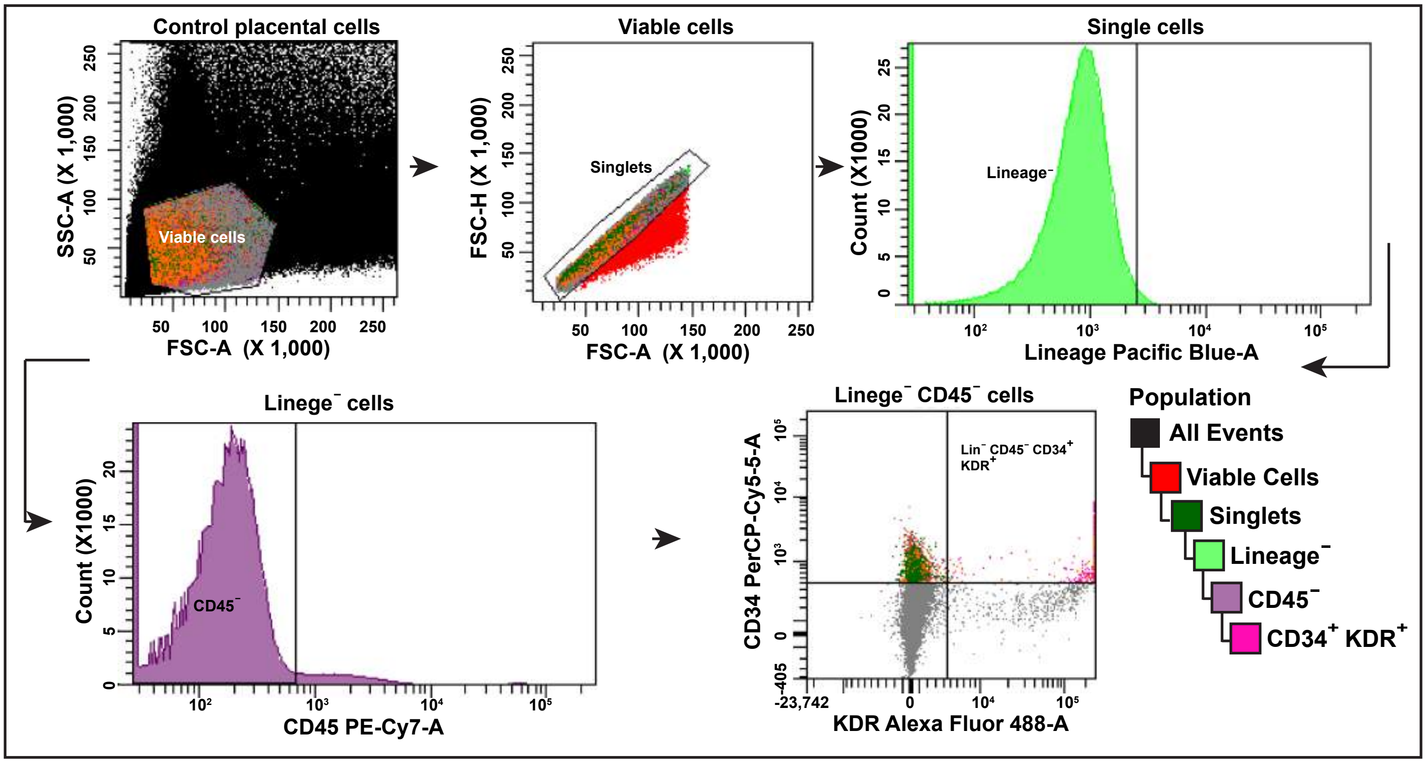


**Fig. S9.** Gating scheme for the flow sort analysis of the $CD34^{+}$ $KDR^{+}$ population in the control and GATA Pl1-KO placentae. The $CD34^{+}$ $KDR^{+}$ placental cells were selected from the $Lin^{-}$ and $CD45^{-}$ population. The quantitative analyses are presented in the Fig. 3F and 3G.

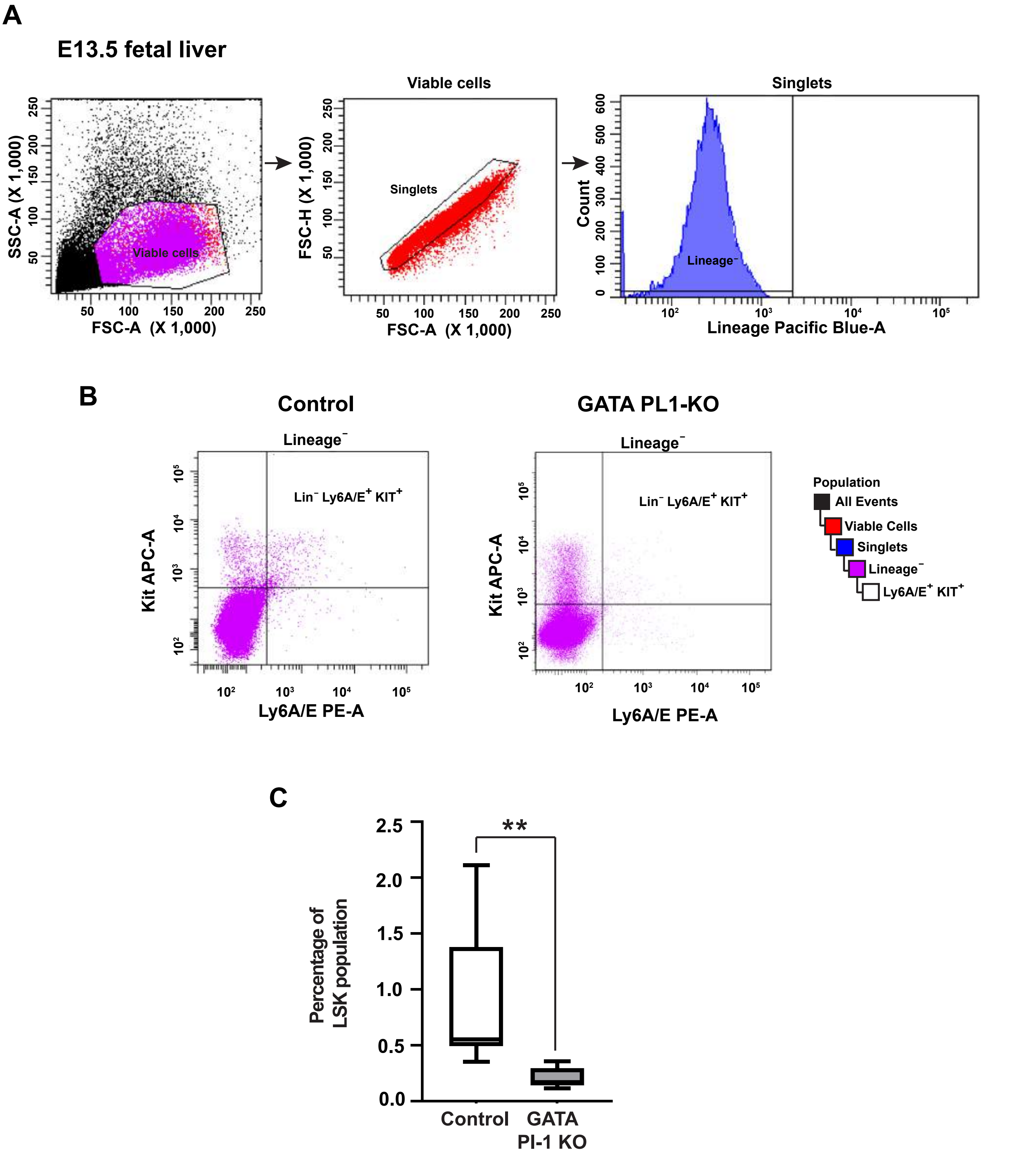


**Fig. S10.** E13.5 fetal liver cell suspensions were subjected to flow analysis. (A) Cells were selected through forward and side scatter followed by singlet selection and Lineage (Lin) negative selection. (B, C) Comparison between the control and the GATA PI1-KO liver samples. Box plot show significant depletion of the Lin$^{-}$ Ly6A/E$^{+}$ KIT$^{+}$ (LSK) population in the knockout population compared to the controls. (Mean±s.em., n=6 for the control and n=12 for the GATA PI1-KO, **P≤0.01, analyzed by two-tailed Student's t-test).

Fig. S11

A

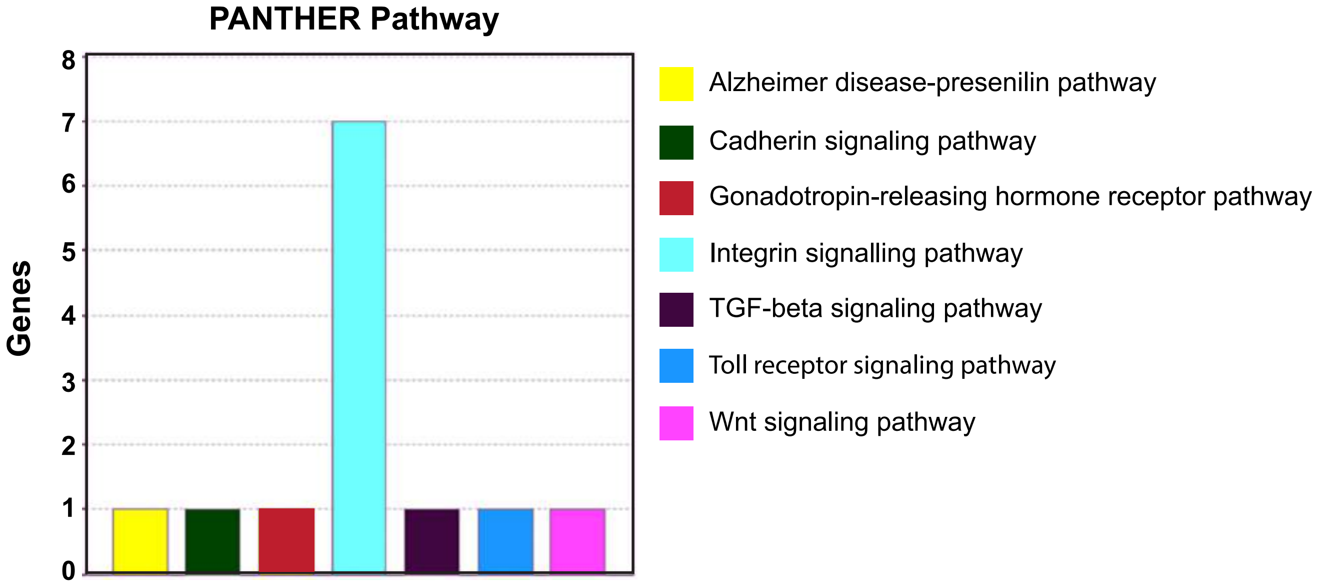


B

| Gene | GATA2 binding | GATA3 binding |
|---|---|---|
| *Col4a1* | Yes | Yes |
| *Col4a2* | Yes | Yes |
| *Fn1* | Yes | No |
| *Inhbb* | No | Yes |
| *Lama5* | Yes | Yes |
| *Lamb1* | Yes | Yes |
| *Lamb2* | Yes | No |
| *Lamc3* | Yes | Yes |
| *Ly96* | Yes | No |
| *Prl2a1* | Yes | No |
| *Prl7d1* | Yes | No |
| *Wnt11* | Yes | No |

C

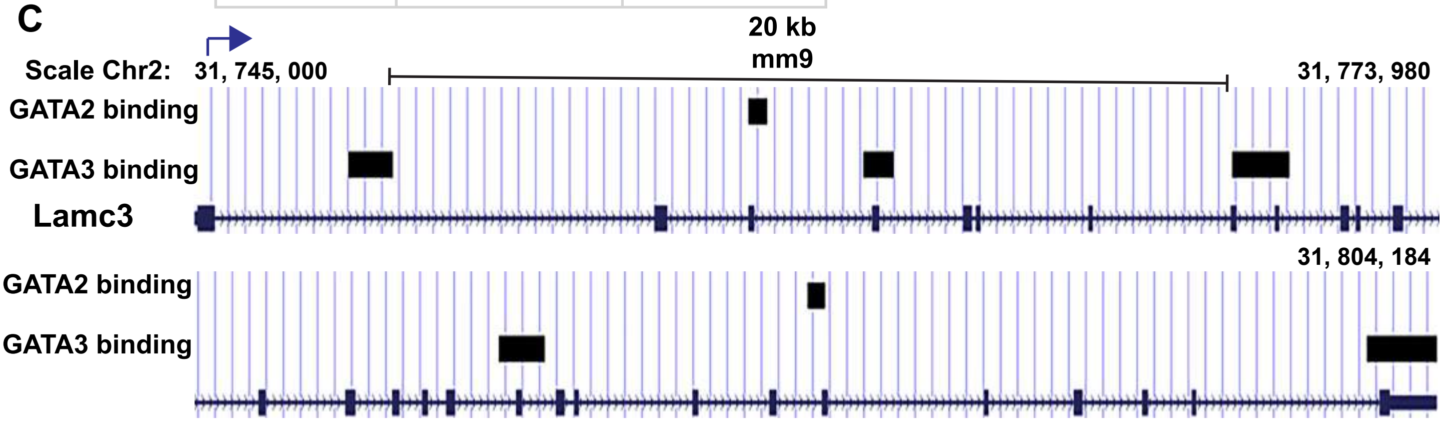


**Fig. S11.** (A) PANTHER pathway analysis of the paracrine factors from *Prl3d1*+ cluster 33 showed 7 members are associated with Integrin signalling pathway. (B) Putative gene targets for GATA2 and GATA3 that are the members of the Integrin signaling pathway. (C) Example of putative binding sites for GATA2 and GATA3 at the *Lamc3* gene locus on chromosome 2.
